# Temperature and Pressure Shaped the Evolution of Antifreeze Proteins in Polar and Deep Sea Zoarcoid Fishes

**DOI:** 10.1101/2024.08.24.609455

**Authors:** Samuel N. Bogan, Nathan Surendran, Scott Hotaling, Thomas Desvignes, Iliana Bista, Luana S.F. Lins, Mari H. Eilertsen, Nathalie R. Le François, Tait Algayer, Scott L. Hamilton, Paul B. Frandsen, Federico G. Hoffmann, Joanna L. Kelley

**Affiliations:** Department of Ecology and Evolutionary Biology, University of California Santa Cruz, Santa Cruz, CA USA; Department of Watershed Sciences, Utah State University, Logan, UT, USA; Institute of Neuroscience, University of Oregon, Eugene, OR, USA; Department of Biology, University of Alabama at Birmingham, Birmingham, AL 35233, USA; LOEWE Centre for Translational Biodiversity Genomics, Frankfurt, 60325, Germany; Senckenberg Research Institute and Natural History Museum, Frankfurt, 60325, Germany; Wellcome Sanger Institute, Tree of Life, Wellcome Genome Campus, Hinxton CB10 1SA, United Kingdom; CSIRO, Health and Biosecurity, Canberra, Australia; Department of Biological Sciences, Centre for Deep Sea Research, University of Bergen, Norway; Laboratoire de Physiologie et Aquaculture de la Conservation (LPAC), Division des collections vivantes, de la conservation et de la recherche, Biodôme de Montréal/ Espace pour la vie, 4777, av. Pierre-De Coubertin, Montréal, Québec H1V 1B3; School of Biological Sciences, Washington State University, Pullman, WA, USA; Moss Landing Marine Laboratories, San Jose State University, Moss Landing, CA, USA; Department of Plant and Wildlife Sciences, Brigham Young University, Provo, UT, USA; Department of Biochemistry, Nutrition, and Health Promotion, Mississippi State University, Mississippi State, MS, USA; Institute for Genomics, Biocomputing and Biotechnology, Mississippi State University, Mississippi State, MS, USA

**Keywords:** Long-read sequencing, antifreeze protein, adaptation, genomics, temperature, gene copy number

## Abstract

Antifreeze proteins (AFPs) have enabled teleost fishes to repeatedly colonize polar seas. Four AFP types have convergently evolved in several fish lineages. AFPs inhibit ice crystal growth and lower cellular freezing point. In lineages with AFPs, species inhabiting colder environments may possess more AFP copies. Elucidating how differences in AFP copy number evolve is challenging due to the genes’ tandem array structure and consequently poor resolution of these repetitive regions. Here we explore the evolution of type III AFPs (AFP III) in the globally distributed suborder Zoarcoidei, leveraging six new long-read genome assemblies. Zoarcoidei has fewer genomic resources relative to other polar fish clades while it is one of the few groups of fishes adapted to both the Arctic and Southern Oceans. Combining these new assemblies with additional long-read genomes available for Zoarcoidei, we conducted a comprehensive phylogenetic test of AFP III evolution and modeled the effects of thermal habitat and depth on AFP III gene family evolution. We confirm a single origin of AFP III via neofunctionalization of the enzyme sialic acid synthase B and show that AFP copies expanded under low temperature, but also decreased with depth, which reduces freezing point via pressure. Associations between the environment and AFP III copy number were driven by duplications of paralogs that were translocated out of the ancestral locus at which Zoarcoidei AFP arose. Our results reveal novel environmental effects on AFP evolution and demonstrate the value of high-quality genomic resources for studying how structural genomic variation shapes convergent adaptation.

## 1. Introduction

The evolution of gene family copy number variation can contribute to environmental adaptation (Schmidt et al. 2010) and be stimulated by environmental change (Hull et al. 2017). Studying the causes and consequences of copy number evolution is difficult when duplications produce large tandem repeats or affect synteny, but the advent of long-read sequencing has helped resolve repetitive variants clouding the origins of copy number evolution (Marx 2023). The evolution of antifreeze proteins (AFPs) and antifreeze glycoproteins (AFGPs) are remarkable examples of convergent environmental adaptation in ectotherms living at low temperatures (DeVries 1971; Baskaran et al. 2021). Increased AFP ‘dosage’ driven by gene duplication can facilitate adaptation to chronic cold (Desjardins et al. 2012). In theory, temperate species derived from cold-adapted ancestors should undergo relaxed purifying selection on AFP coding regions, relaxed selection on AFP copy number, and experience subsequent AFP copy number contractions (Hew et al. 1988; Hayes et al. 1991). Studying contractions and expansions of AFPs across thermally-distributed clades is critical for understanding thermal adaptation by ectotherms but has been hampered by poor resolution of their copy number due to low quality reference genome assemblies.

Antifreeze proteins were first discovered in Antarctic notothenioid fishes, which possess antifreeze glycoproteins (AFGPs). AFGPs were found to lower both the freezing and boiling point of cellular fluid (DeVries and Wohlschlag 1969; DeVries 1971; Raymond and DeVries 1977). Since this discovery, AFPs that induce thermal hysteresis, an increase in degrees of difference between a fluid’s freezing and boiling points (Barrett 2001), have been discovered in microbes (Gilbert et al. 2005), plants (Gupta and Deswal 2014), invertebrates (Duman 2015), and vertebrates (DeVries and Cheng 2005). AFPs generally function by binding to the leading edges of newly-crystalized nuclei of ice, attenuating their expansion and thereby reducing cellular freezing point (Jia et al. 1996). In fishes, AFPs have independently evolved at least four times (Cheng and Devries 2002; Davies 2022). These convergently evolved AFPs make up four known categories termed AFGPs, and AFP types I, II, and III. The evolution of AFPs by ectotherms is a classic example of environmental adaptation, but we still know little about the environmental processes that shaped their evolution and the diversity and variation they exhibit across species and their genomes.

It has been repeatedly suggested that, among lineages possessing AFPs, species or populations inhabiting colder environments should exhibit more AFP copies (Hew et al. 1988; Hayes et al. 1991; Kelley et al. 2010). Comparative studies of AFP copy number in population– and species-pairs of fishes from nominally warmer versus colder habitats have confirmed this expectation (Hayes et al. 1991; Desjardins et al. 2012). Prior research demonstrates that AFP genes are lost in warmer, non-freezing habitats. Studying AFGPs in Arctic cod and Antarctic notothenioids, Baalsrud et al. (2018) observed losses of AFGPs in secondarily sub-polar species of both groups. In addition to evolving neutrally above this threshold, AFPs may be subject to negative selection suggestive of costs to AFP activity. Indeed, the expression of AFPs and AFGPs can disrupt organismal function through accumulation of superheated ice crystals (Celik et al. 2010; Cziko et al. 2014) and potentially impose costs of AFP activity in individuals experiencing fluctuating freezing risk, such as many Arctic fishes (Scholander et al. 1953). Long-read sequencing of notothenioids also revealed pseudogenization and contraction of AFGP copies in individual species inhabiting milder latitudes suggestive of relaxed purifying selection on AFP sequences or negative selection on antifreeze function (Bista et al. 2023; Rivera-Colón et al. 2023). However, an analysis of AFP copy number across continuous thermal clines has not been reported.

Whether and how AFP copy number and gene dosage scales with freezing risk remains unknown. One environmental variable that has been proposed to affect freezing risk and AFP evolution is pressure. Because increased pressure in seawater affects the stability of ice crystals and decreases phase change temperatures, thus lowering freezing point (Fujino et al. 1974), it has been proposed that polar fishes inhabiting different depths and exhibiting divergence in AFP copy number have experienced differential purifying selection or differential directional selection on AFP copy number and activity (Desjardins et al. 2012). Deep ocean fishes also regulate hydrostatic pressure by increasing cellular and serum osmolite content (Yancey et al. 2002; Yancey et al. 2014), which may further reduce internal organismal freezing temperatures.

AFPs of bony fishes have evolved by neofunctionalization, and AFGPs by both neofunctionalization and *de novo* gene birth. These events were driven in part by repetitive genomic regions that provided substrate for *de novo* gene birth of an AFGP in cods (Baalsrud et al. 2018; Zhuang et al. 2019) and AFP I flounder (Rives et al. 2024). AFP III evolved in Zoarcoidei fishes via translocation of ancestral genes that underwent neofunctionalization and expanded into tandem arrays of AFPs (Deng et al. 2010; Hotaling et al. 2023). AFP I also evolved via neofunctionalization in sculpins (Graham and Davies 2024). In addition to influencing AFP origins, structural variation such as translocations, inversions, and repetitive element content have the potential to influence expansions and contractions of AFPs across environmental gradients (Lauer and Gresham 2019). For example, cytogenetically-balanced and unbalanced translocations can both facilitate variation in gene copy number (Watson et al. 2007; Harewood et al. 2010). The accurate detection of translocations and measurement of gene copy number in repetitive regions require long-read sequencing approaches, which enable identification of structural variants such as translocations and better resolve complex genomic regions relative to short read sequencing (Marx 2023). Long-read sequencing has shown success in assembling repetitive AFGP arrays in notothenioid fishes (Bista et al. 2023; Rivera-Colón et al. 2023)

The bony fish suborder Zoarcoidei is an ideal system in which to study the joint effects of environmental and structural genomic variation on AFP evolution. Zoarcoidei likely originated in north temperate seas, followed by stepwise invasions of (i) the Arctic Ocean, (ii) south temperate seas and (iii) the Southern Ocean (Anderson 1994; Radchenko 2016; Hobbs et al. 2020; Hotaling et al. 2021) where minimum temperatures can fall below –2.0 ℃ (Fetterer 2017; Auger et al. 2021). Multiple deep sea environments exhibit temperatures below 0 ℃ and as low as –2.0 ℃ (Ponte 2013). These extremely low temperatures can cause freezing in tissues and serum (Storey and Storey 1988) and the formation of sea ice at ocean surfaces and as brinicles within the water column (Worster and Rees Jones 2015). Fish tissues typically possess higher freezing points than seawater as they are hypoosmotic to their environment (Hargens 1972), posing freezing risk even in the absence of sea ice. Sanger sequencing of BAC clones in the zoarcoid *Lycodichthys dearborni* suggested that type III AFP arose within Zoarcoidei via neofunctionalization of the enzyme sialic acid synthase B (*sasb*) (Deng et al. 2010). *sasb* likely duplicated before undergoing deletions of all exons except 1 and 6, which are homologous to AFP III exons 1 and 2 (Deng et al. 2010). Zoarcoidei repeatedly invaded and adapted to the poles (Hotaling et al. 2021) while Zoarcoidei AFP III likely evolved once (Deng et al. 2010; Hobbs et al. 2020).

AFP III copies have likely expanded and contracted multiple times in Zoarcoidei (Hotaling et al. 2023), but the extent of gene family evolution of AFP III in zoarcoids, of AFPs across fishes, and how genomic and environmental variation interactively affected AFP evolution remain unclear. Zoarcoidei diversified across depth faster than any group of fishes (Friedman and Muñoz 2023), potentiating a complex landscape of selection for AFP function and dosage across the suborder’s distributions through latitude and depth. Previously identified tandem arrays of zoarcoid AFP III exhibited conserved synteny with several flanking genes (Hotaling et al. 2023), enabling identification of translocations out of this region.

Here we report high-quality, long-read genome assemblies from six species of Zoarcoidei from diverse thermal habitats and depths (Fig. 1; Table 1). We integrated annotations of AFP III genes with oceanographic measures of thermal minima, pressure, and depth within species’ habitats to test the hypothesis that temperature and pressure are associated with AFP III copy number evolution. We predict that AFP copies contracted in deep ocean, higher-temperature species through relaxed purifying selection or negative directional selection on AFP function and copy number. Using analyses of synteny, we traced AFP III copies to ancestral and translocated arrays and determined whether translocation drove AFP expansions under high freezing risk. These new reference genome assemblies provide a valuable resource for studying how abiotic environments and structural variation jointly influence convergent adaptation.

**Figure 1.**
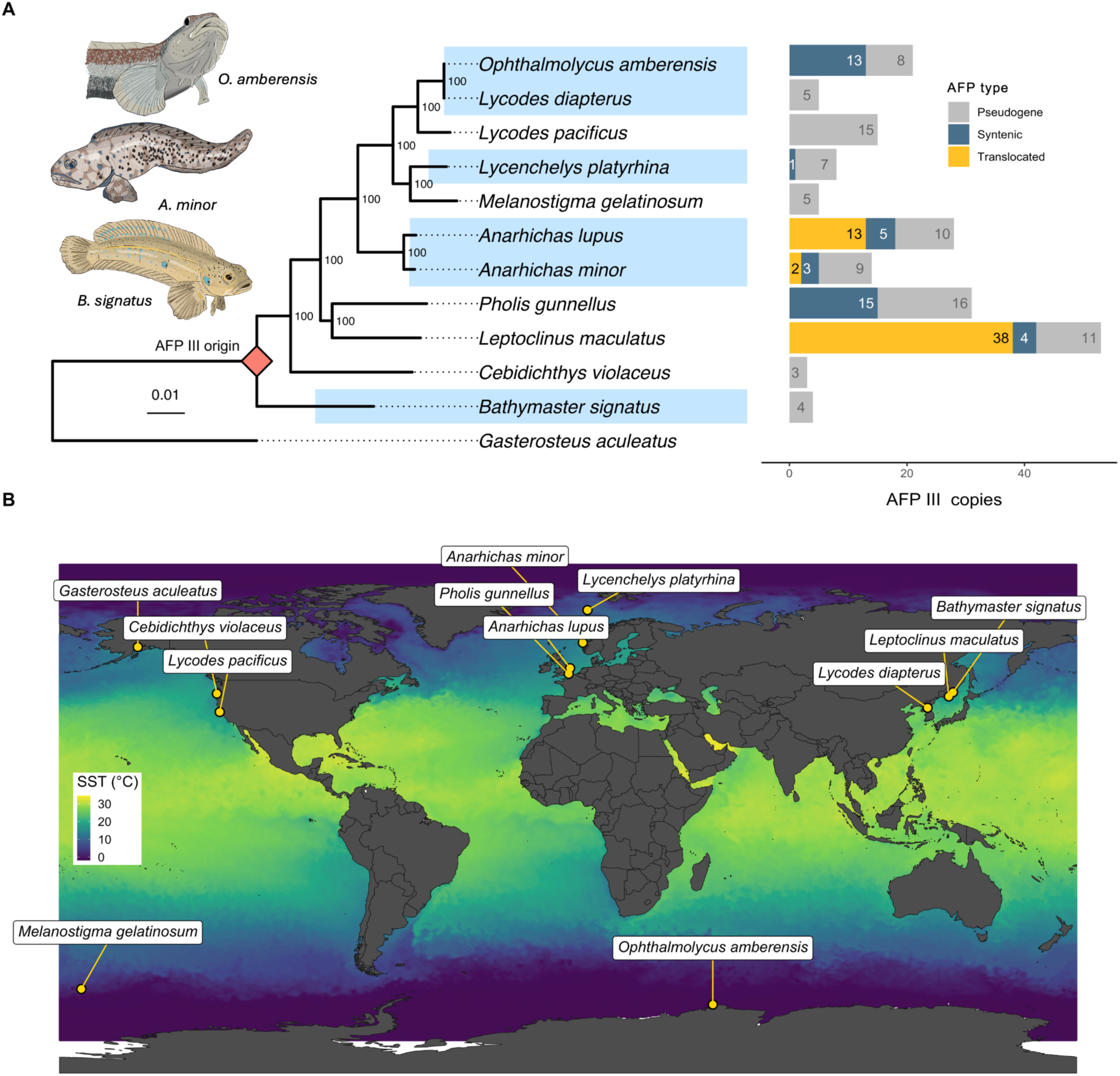
| Species’ phylogeny, AFP copy numbers, and distributions. (A) A species tree of Zoarcoidei included in this study including outgroup *Gasterosteus aculeatus* is plotted adjacent to AFP copy number variation colored by pseudogenization or location of homologs in the ancestral syntenic region or translocation(s). The red diamond signifies the origin of type III AFP. Nodes are labeled with bootstrapped support values. Species highlighted in blue denote genome assemblies reported by the authors in this study and Hotaling et al. (2023). The scale bar denotes substitutions per amino acid. *Ophthalmolycus amberensis* and *Lycodes diapterus* branch tip lengths equal 0.000069 and 0.000074 substitutions per amino acid, respectively. (B) Species’ distributions are plotted as the point of maximum likelihood of occurrence from AquaMaps standardized distributions. The size of white circles around occurrences are scaled according to species abundance. Mean sea surface temperature (SST, °C) recorded in August, 2020, is visualized according to color. Illustrations by SNB.

**Table 1.**
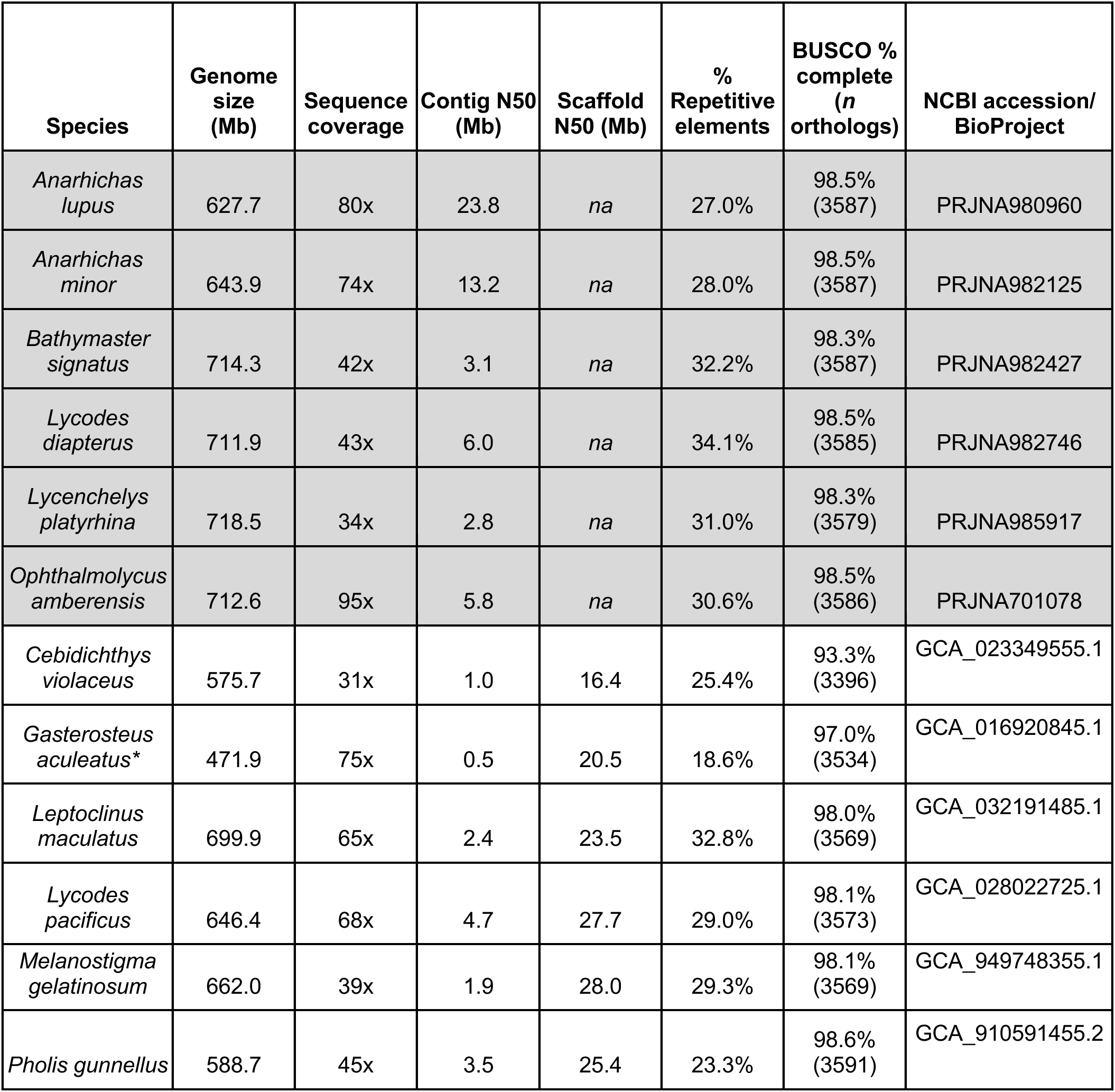
| Genome assembly metrics. Genomes include those reported by the authors in this study (rows 1-5) and Hotaling et al. 2023 (row 6). Assemblies reported by the authors are shaded in gray. Publicly available zoarcoid genomes leveraged in this study are not highlighted. A non-zoarcoid outgroup is designated by an asterisk. BUSCO analysis was performed on universal single-copy orthologs of Actinopterygii.

## 2. Results and Discussion

### 2.1. Genome assembly

Reference genomes were assembled with Pacific Biosciences (PacBio) long-read sequence data for six zoarcoid fishes spanning a range of temperatures and depths: the ronquil *Bathymaster signatus*, the wolffishes *Anarhichas lupus* and *Anarhichas minor*, and the eelpouts *Lycenchelys platyrhina*, *Lycodes diapterus*, and *Ophthalmolycus amberensis*. These species inhabit polar seas with latitudes as northerly as 73.0 °N and southerly as –65.8 °S. Assembled genomes included temperate species with absolute latitudes as low as 37.8 ° (Fig. 1). Species’ maximum depths spanned 24 – 2,229 meters. These mark the first long-read genome assemblies for the families Bathymasteridae (ronquils) and Anarhichadidae (wolffishes), and increase the number of long-read genome assemblies among Zoarcidae (eelpouts) by two and a half times.

Genome sizes, sequencing coverage, N50 values, BUSCO complete percentages, and percent repetitive elements are reported in Table 1 for the six new assemblies, five existing zoarcoid long-read assemblies (Rhie et al. 2021; Programme et al. 2022; Wright et al. 2023; Bista et al. 2024), and one scorpaeniform outgroup (Nath et al. 2021). Genome sizes of all Zoarcoidei examined equaled a mean of 647.80 ± 74.3 Mb. Genome assemblies reported here averaged 11.7x greater coverage compared to publicly available, long-read zoarcoid assemblies. Contig N50 values were an average of 6.5 Mb longer than available assemblies (Table 1). As expected, repetitive element content linearly and positively correlated with genome size across all species (Kidwell 2002) as shown in Fig. S1 (r^2^ = 0.93).

Zoarcoidei families are among the few fish groups inhabiting both the Arctic and Southern Oceans (Møller et al. 2005), positioning this suborder to be a valuable system in which to study convergent evolution and adaptation to extreme cold. However, a paucity of genomic resources for zoarcoid fishes stands as an obstacle to their value in research. The improved reference genome quality of these new assemblies among other members of Zoarcoidei and the extension of thermal habitats and depths represented among them significantly strengthen Zoarcoidei as a tractable, valuable system in which to study convergent adaptation.

### 2.2. Variation in type III antifreeze protein copy number

The number of AFP III copies in the 11 zoarcoids studied ranged from 0 to 42 putatively functional copies. Between 3 and 16 pseudogenized AFP III copies were detected across all zoarcoids (Fig. 1A). The copy number of putatively-functional and pseudogenized AFP genes correlated such that pseudogenes increased in species with more AFP III copies (Fig. S2; r^2^ = 0.31). This association was explained by phylogeny and became insignificant after controlling for species relatedness (95% posterior interval = –1.54 – 0.67). The presence of pseudogenized AFP III copies in all zoarcoids indicates that the gene’s origin predates the most-recent common ancestor of zoarcoids included in this study (Fig. 1A). From here forward, all results correspond to the study of putatively functional AFP copies and do not include pseudogenes.

We evaluated whether AFP III may have arisen in a common ancestor predating Zoarcoidei by screening for the gene in long-read assemblies of 12 species of the Order Scorpaeniformes, representing 7 families, and found no evidence of complete or partial AFP III paralogs (Table S2). Past phylogenetic analyses of zoarcoid AFP III also detected a partial, pseudogenized AFP III sequence in Bathymasteridae, the earliest-branching family of Zoarcoidei (Hobbs et al. 2020). However, Hobbs et al. (2020) and others have not ruled out an emergence of AFP III prior to the branching of Zoarcoidei. Our results therefore strengthen evidence that Zoarcoidei AFP III likely arose in a common ancestor Bathymasteridae and Zoarcoidei as opposed to a common ancestor within Scorpaeniformes predating Zoarcoidei. Long-read sequencing of Zoarcoidei offers the potential to trace the origin of AFP III within Zoarcoidei by mapping structural variants contributing to the gene’s evolution. These events include duplication-translocation of *sasb* and deletions of *sasb* exons 2 – 5.

### 2.3. Environmental effects on type III antifreeze protein copy number

Habitat temperature and pressure jointly influence freezing risk (Fujino et al. 1974) and were additively correlated with AFP copy number such that AFP copy number expanded in shallow, cold-water species (Fig. 2A). A phylogenetically corrected Bayesian model revealed that AFP copy number significantly increased with decreases in (i) thermal minima of species’ habitats and (ii) mean pressure of species’ depth ranges (Fig. 2). Thermal minima equaled the lowest recorded temperature at a species’ coordinate of maximum likelihood of occurrence within their depth range. Pressure was derived from habitat depth and latitude. Thermal minima of species habitats spanned –3.00 to 7.21°C (Fig. S3). AFP copy number significantly increased in species occupying lower thermal minima (Fig. 2B). This effect was significant after controlling for phylogeny (scaled effect = –1.55 ± 0.97). It is possible that a thermal gradient in AFP III copy number evolved in Zoarcoidei via mutation that preceded and enabled invasion of novel temperatures or changes in copy number that followed the invasion of new temperatures inducing selective pressure.

**Figure 2.**
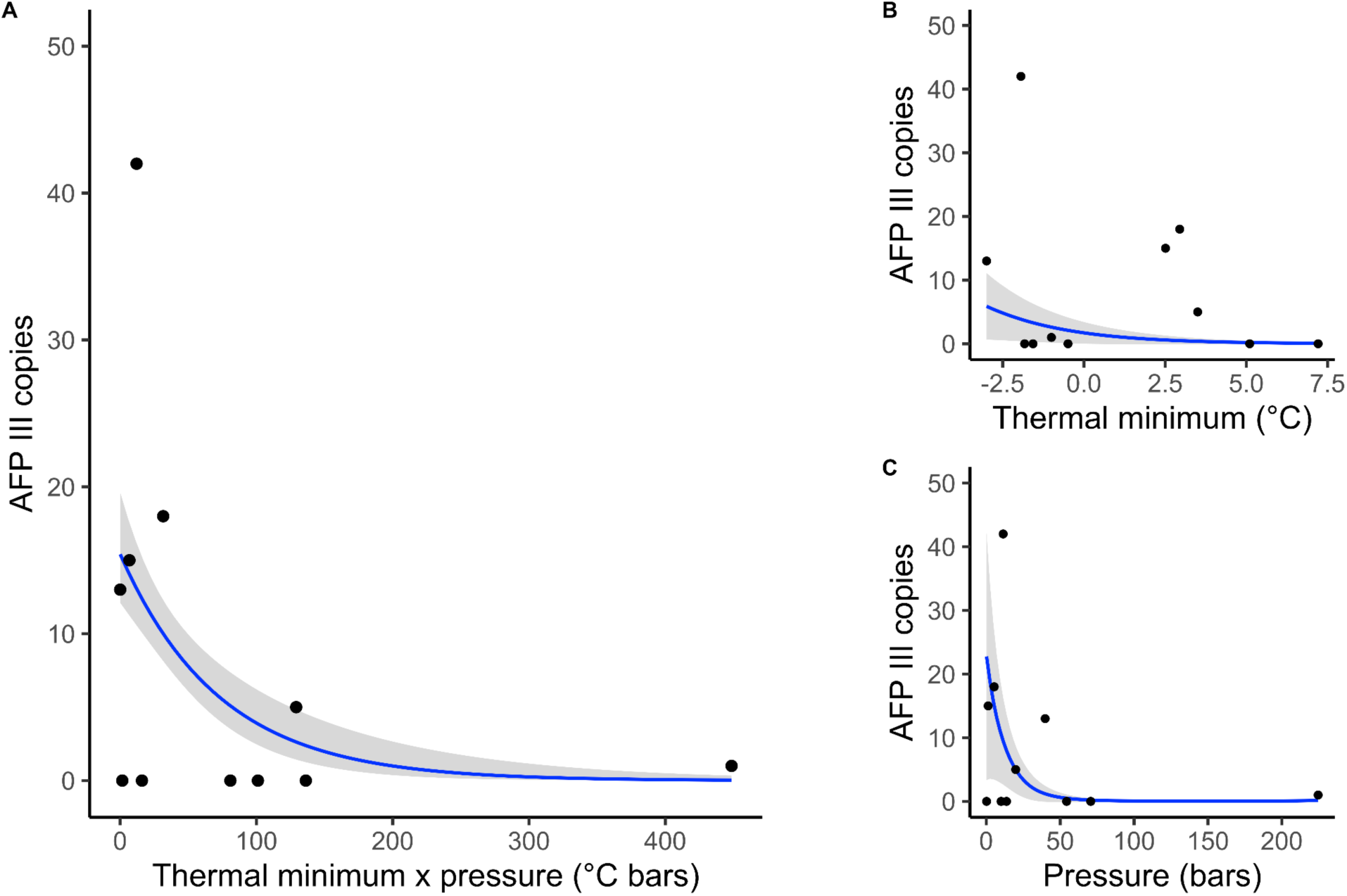
| Associations between temperature, pressure, and type III antifreeze protein copy number. (A) AFP expansions in shallow, cold-water species are visualized as a logistic regression of AFP copy number fitted to the product of mean habitat pressure and habitat thermal minimum, with the lowest thermal minimum scaled to 0. Panels on the right show relationships between AFP III copy number and (B) thermal minima of species habitats and (C) mean pressure of species’ habitats are plotted. Fitted lines represent fit from a phylogenetically controlled model of type III antifreeze protein copy number.

AFP III gene copy number declined in species inhabiting greater pressure (Fig. 2C). Pressure had a stronger effect on AFP copy number than temperature and was non-linear, following a negative exponential curve. Pressure, which varies as a function of depth and latitude, affected AFP copy number after controlling for habitat depth as well as phylogeny (scaled linear effect = –4.72 ± 2.58). This suggests that pressure, rather than depth, predicted AFP copy number. Habitat temperatures correlated with depth and pressure (Fig. S4 – S5). To account for this autocorrelation, predicted effects of thermal minima and pressure on AFP copy number were modeled in a statistically independent manner. The finding that AFP III copy number is reduced in deep sea species experiencing high pressure is pertinent to the biogeography of Zoarcoidei, which includes the eelpout family Zoarcidae that has exhibited faster rates of evolution across depth than any group of fishes (Friedman and Muñoz 2023). Associations between depth and AFP copy number were previously proposed by Desjardins et al. 2012, who observed that the wolffishes *A. lupus* and *A. minor* exhibit higher AFP copy number in the shallow water species *A. lupus* rather than *A. minor* the species occupying a colder thermal environment (Desjardins et al. 2012). Our results mark the first quantitative evidence that selection on antifreeze proteins across thermal environments was shaped by pressure.

A direct explanation of the association between pressure and AFP copy number is that increased energy imposed by pressure lowers the freezing point of intracellular fluids (Fujino et al. 1974). An indirect explanation also exists that is not mutually exclusive to this direct hypothesis. Deep sea fishes generally regulate the hydrostatic pressure of cells through osmoregulation, such that increased pressure is countered by greater osmolarity in cells and serum, where AFPs are secreted (Griffith 1981; Gillett et al. 1997; Yancey et al. 2002; Yancey et al. 2014). Because increased osmolality also reduces the freezing point of fluids (Potter et al. 1978; Bodnar 1993), it pressure and osmolality may have additively affected freezing point.

It is possible that reductions in AFP gene copy number in species of warmer minimum habitat temperature can be attributed to relaxed purifying selection on AFP sequence and/or negative selection on AFP copy number or AFP function. Many polar fishes inhabiting the Arctic Ocean and almost all polar fishes included in our study, excluding *O. amberensis,* experience temporal and spatial variation in freezing risk across their range and throughout the year (Reist et al. 2006; Verde et al. 2006). It is likely that fishes which have expressed AFPs to attenuate ice growth under freezing conditions incur a cost to AFP function during episodic non-freezing conditions. Fishes that experience more frequent temperatures above freezing point may experience stronger negative selection on AFP expression level or function, which would select for reduced copy number and/or AFP activity. Additionally, some non-polar populations share connectivity with polar populations experiencing freezing risk (Hickerson and Cunningham 2006) and populations that do not experience freezing risk may possess allele frequencies that have been shaped by selection on AFP sequence or function in connected, polar populations.

### 2.4. Gene family evolution and convergence of antifreeze protein copy number

Restricted maximum likelihood ancestral state reconstruction of AFP III copy number demonstrated that at least 8 expansions and 12 contractions of AFP III occurred across the studied phylogeny (Fig. S6). To evaluate whether AFP contractions and expansions convergently evolved during independent invasions of polar or deep seas, the phylogenetically-controlled model of environmental associations with AFP III copy number was expanded as a structural equation model predicting both habitat thermal minima and mean pressure as a function of phylogenetic relatedness. This enabled regression of AFP III copy number against phylogenetically-independent invasions of colder or higher-pressure environments, which is necessary for quantifying convergence. Independent invasions of lower thermal minima were significantly associated with increased AFP III copy number (95% posterior interval = –1.15 – –0.03 AFP copies per °C). Phylogenetically-independent changes in habitat thermal minima and AFP copy number are plotted in Figures S7-8. Despite having a stronger association with pressure before testing for convergence, independent invasions of higher pressure habitats were not associated with decreased AFP III copy number.

While the association between AFP III copy number and habitat pressure among extant branches was stronger than with thermal minimum, there is little evidence that this association resulted from convergent evolution between independent invasions of the deep ocean. By contrast, weaker associations between temperature and AFP III are more likely to have evolved as a result of convergent gains and losses of AFP III during invasions of Zoarcoidei into colder and warmer environments, respectively. The pole-to-pole distribution of Zoarcoidei is marked by a likely origin in the Arctic followed by invasion of the deep ocean, trans-polar migration across deep-ocean regions, and subsequent reinvasion of shallow waters in the Southern Ocean (Hobbs et al. 2020; Hotaling et al. 2023). The ancestral state reconstruction of thermal minima shows repeated invasions of sub-zero oceans across Zoarcoidei (Fig. S7). As shown in Figure S9, the reconstruction of habitat pressure shows one primary lineage of deep ocean fishes belonging to the eelpout family Zoarcidae, which is the deepest dwelling family of zoarcoid fishes. Thus, the strong effect of pressure on AFP III copies across Zoarcoidei was more likely attributed to a singular invasion of the deep sea by Zoarcidae rather than multiple convergent events. Decreases in habitat pressure in Zoarcidae that corresponded with increased AFP III copy number represent instances of parallel rather than convergent evolution. Phylogenetically-independent changes in habitat pressure and AFP copy number are plotted in Figures S9-10.

Using maximum likelihood gene tree inference, we confirmed the origin of AFP III evolution in Zoarcoidei. Our tree of all *sasa*, *sasb*, and AFP III homologs across study species is consistent with a single origin of AFP III from *sasb* (Deng et al. 2010) as shown in Figure 3C. However, the topology of the gene tree does not provide evidence of whether the common ancestor of Zoarcoidei arose at the syntenically-conserved array or translocated locus. For the gene tree to illuminate the positional origin of Zoarcoidei AFP, a monophyletic cluster of syntenic or translocated AFPs would have to be the first lineage to diverge from all other AFPs. The gene tree does not show a monophyletic cluster of translocated AFPs as the first diverging group of Zoarcoidei AFP. Therefore, maximum likelihood ancestral state reconstructions were used to predict binary characters of whether species possessed AFP III in the syntenic locus or in one of the six translocated loci. These reconstructions demonstrated that ancestral Zoarcoidei AFP III most likely arose in the syntenic region rather than one of the translocated positions (relative likelihood = 727.89 – 1440.39).

**Figure 3.**
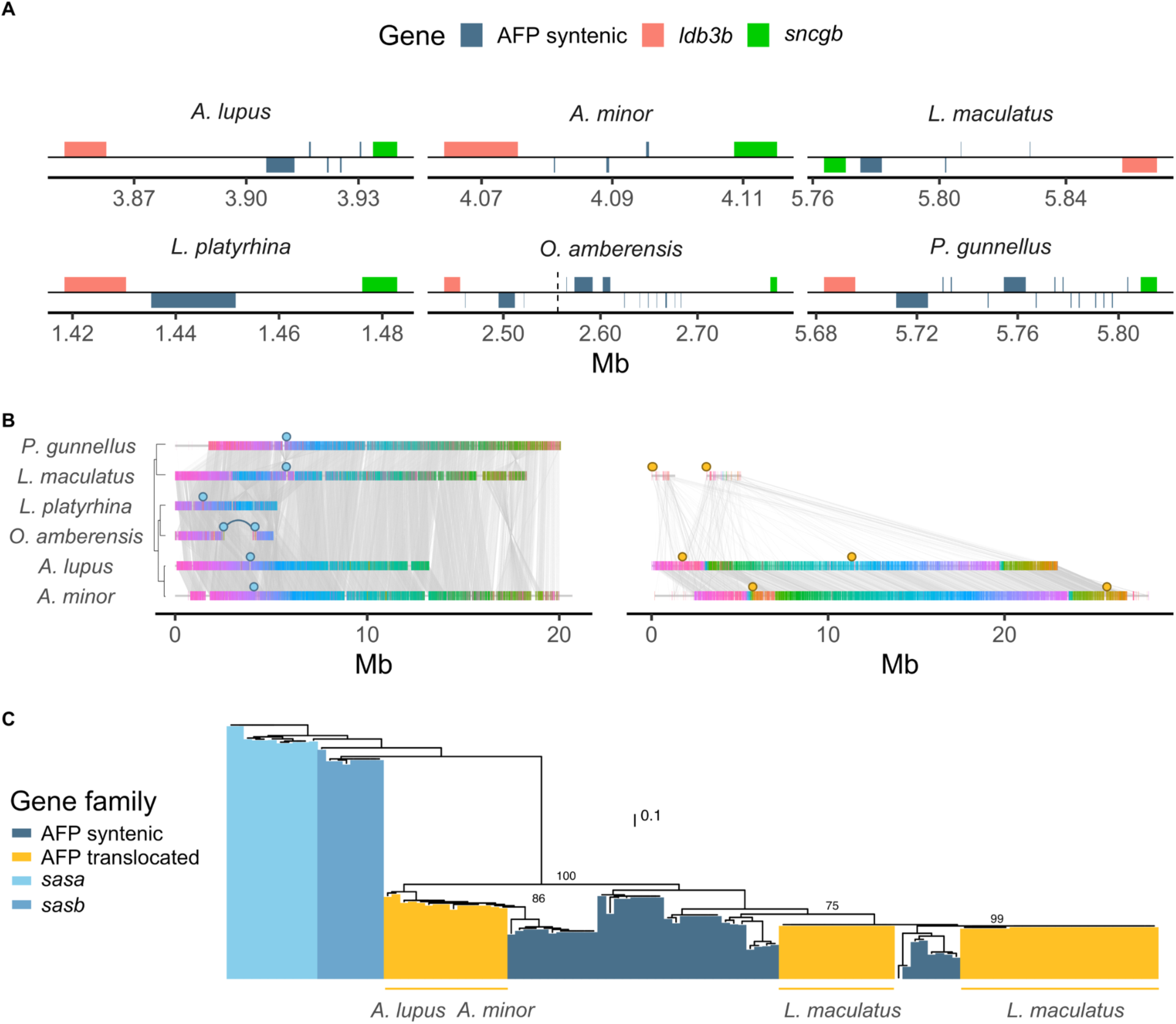
| Synteny and origin of type III antifreeze proteins. (A) Microsynteny at the ancestral, syntenic Zoarcoidei AFP array is visualized by annotation tracks colored by whether they represent AFPs or genes flanking the array (*ldb3b* and *sncgb*). The vertical dashed line in *O. amberensis* represents a junction between contig fragments. Tracks represent the full lengths of gene exons and introns. (B) Synteny of AFP III tandem arrays. Full lengths of scaffolds and contigs are shown. Colored bars represent regions conserved in at least two species. Regions sharing the same color are homologous. Gray lines connect these homologous regions. Narrow horizontal bars depict regions lacking homology. Positions of ancestral, syntenic AFP arrays are marked by blue circles and translocated arrays by orange circles. Curved lines connect single AFP arrays split between contigs. A species distance tree is inset into the left hand side of the syntenic array panel. (C) A rooted gene tree of *sasa*, *sasb* and AFP III paralogs. Branches are colored by syntenic AFPs, translocated AFPs, *sasa*, or *sasb* genes. The scale bar represents nucleotide substitutions per base pair. Bootstrap support values are included to the right of all nodes that branch into clades of syntenic and translocated AFPs. The taxonomic groups that translocated AFPs belong to are annotated below translocated AFP clades. The node most closely related to *sasb* branches into gene clades of translocated and syntenic AFPs, prohibiting resolution of whether ancestral Zoarcoidei AFP evolved in the syntenically conserved region. Tests of whether Zoaroidei AFP arose in the syntenic region are described under ‘*Gene family evolution and convergence of antifreeze protein copy number*’.

### 2.5. Role of translocation in AFP III expansions

The ancestral AFP III array falls within a conserved, syntenic region flanked by genes including synuclein gamma (*sncgb*) and LIM-binding domain protein (*ldb3b*) as previously shown by Deng et al. (2010). We identified at least six independent translocations of AFP III out of the ancestral syntenic region (Fig. 3B). All zoarcoids with AFP III retained this microsynteny, such that at least some AFP III copies were flanked upstream by *sncgb* and downstream by *ldb3b* (Fig. 3A). Whole-contig alignment revealed conservation of synteny across larger genomic scales. The ancestral AFP III tandem repeat region, flanked by *sncgb* and *ldb3b*, fell in the same homologous region of chromosome 21 across Zoarcoidei (Fig. 3B). In *L. maculatus*, however, the ancestral AFP III array was positioned in a ∼3 Mb chromosomal inversion (Fig. 3B). Translocated AFPs clustered into three gene clades (Fig. 3C). In *P. gunnellus*, one AFP copy on chromosome 21 fell outside of the flanking genes *sncgb* and *ldb3b* (Fig. 3A).

AFP III copies outside of the ancestral array mapped to two translocated regions in each of three species, totalling six translocated arrays across all study species. The wolffishes *A. lupus* and *A. minor* each contained two translocations on contigs homologous to *P. gunnellus* chromosome 2. However, the two translocated arrays in *A. lupus* and the two in *A. minor* possessed differing synteny, falling into four distinctly different regions of chromosome 2 (Fig. 3B). Thus, these arrays likely represent four discrete translocation-duplication events. The two translocated arrays in *L. maculatus* were distributed across two contigs mapping to different regions of *P. gunnellus* chromosome 12 (Fig. 3B). Translocated copies contributed significantly to AFP III expansions across Zoarcoidei, making up 13 of 18 (72%) AFP copies in *A. lupus*, 2 of 5 (40%) copies in *A. minor*, and 38 of 42 (90%) copies in *L. maculatus* (Fig. 1A). Considering the short lengths of contigs in which translocated AFP III were located *L. maculatus* (Fig. 3B), there is potential for misassembly to have inflated the total number of translocations, the AFP copy number within translocated arrays, and the total AFP copy number in this species. Conversely, translocated arrays that border the ends of these contigs may also cause underreporting of total AFP copy number at these locations (Fig. 3B).

A multivariate phylogenetic model predicting syntenic and translocated AFP III copy number as a function of thermal minima and pressure revealed that environmental gradients in AFP copy number were driven by expansions of translocated homologs. After controlling for phylogeny, habitat thermal minima was associated with syntenic AFP copies at a scaled effect strength of –0.62 ± 0.72 and translocated AFP copies at –1.36 ± 0.70 (Fig. 4A). Mean habitat pressure yielded an effect on syntenic copies of –0.88 ± 0.54 and an effect on translocated copies of –9.84 ± 3.08 (Fig. 4B).

**Figure 4.**
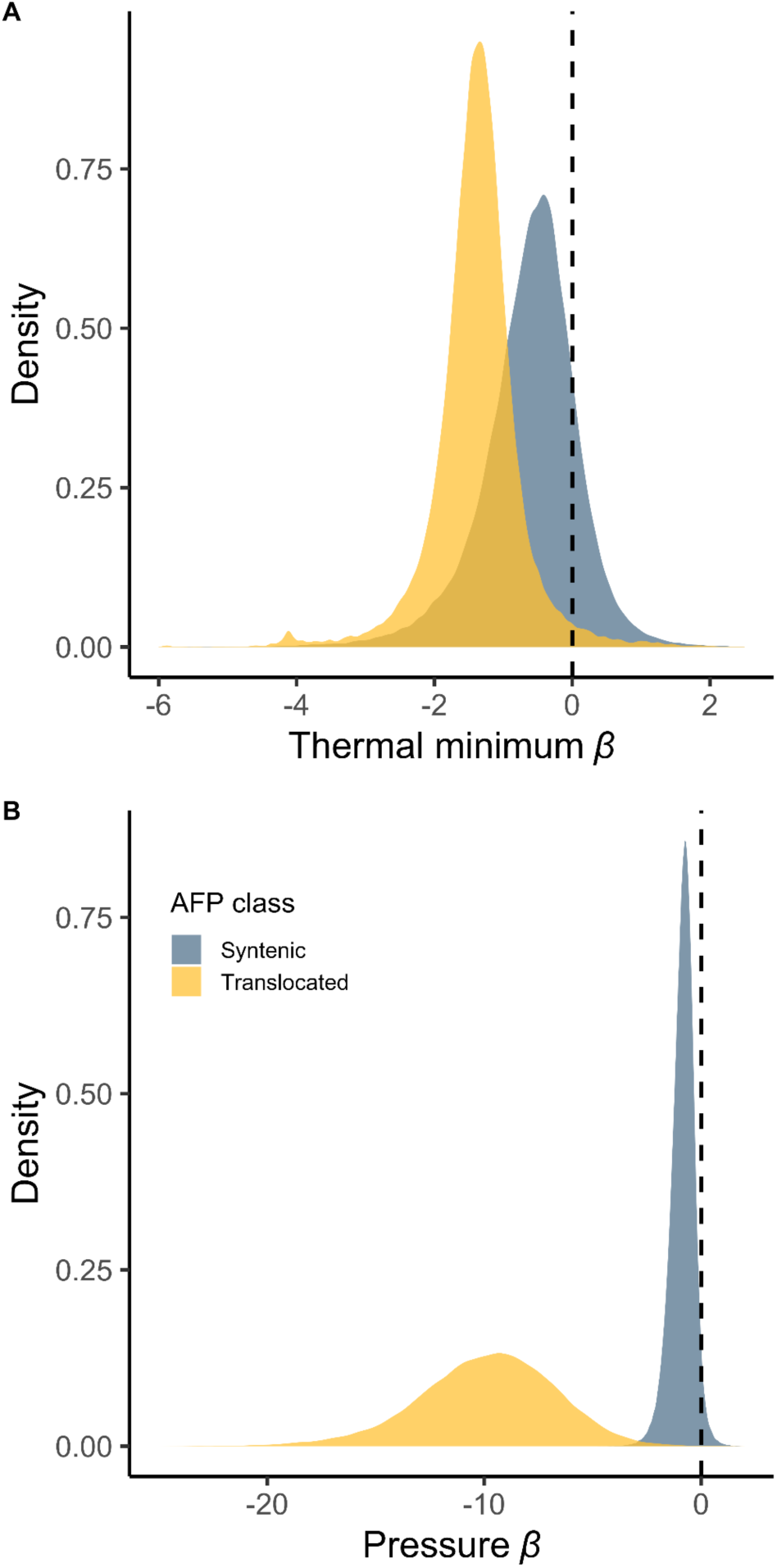
| Effects of thermal minimum and pressure on copy number of syntenicically-conserved and translocated type III antifreeze protein genes. Densities represent the posterior probabilities for fixed effects (β) of (A) thermal minimum and (B) mean pressure on antifreeze protein copy in ancestral, syntenically-conserved versus translocated sites.

Molecular repair mechanisms causing variation in copy number such as break-induced replication can directly facilitate translocation of gene copies to regions sharing microhomology with break sites (Hastings, Lupski, et al. 2009; Hastings, Ira, et al. 2009; Kramara et al. 2018). It is possible that expansions of AFP III via translocation are simply a consequence of selection for high copy number. AFP III translocations were not more likely among species with a large number of AFPs, however. Rather, copy number in translocated arrays correlated with habitat temperature and pressure. This is suggestive of directional selection acting more strongly on AFP copy number outside of the syntenic, ancestral AFP III locus. Differences in directional selection on copies may arise as a consequence of subfunctionalization stemming from transcriptional changes that occur following translocation into new regulatory landscapes (Harewood et al. 2010; Wang et al. 2012). Alternatively, copy number at the ancestral AFP array may be under stronger stabilizing selection and evolutionary constraint relative to translocated arrays.

Bayesian Ornstein-Uhlenbeck phylogenetic mixed models were used to evaluate copy number of syntenic and translocated AFPs across the Zoarcoidei tree. We observed 7.03x greater stabilizing selection (α) on syntenic copy number relative to translocations. Differences in selection were significant as evidenced by a Bayes factor test of α posterior probabilities yielding BF = 41.77 (Fig. 5A). Purifying and diversifying selection on coding sequences were modeled across the AFP III gene tree using branch-tip estimates of *d*_N_/*d*_S_. Syntenic and translocated AFPs were both under strong purifying selection, demonstrating that translocated AFPs remain evolutionarily conserved. Translocated AFPs exhibited lower *d*_N_/*d*_S_ ratios than syntenic copies, but this difference was insignificant. Genes with longer branch lengths had significantly higher *d*_N_/*d*_S_ ratios on average as shown in Figure 5B (scaled posterior interval = 0.17 – 0.89). It is possible that translocated AFPs exhibited stronger purifying selection in our test, not because they have truly been under stronger purifying selection, but because of an analytical artifact that stems from having duplicated more recently.

**Figure 5.**
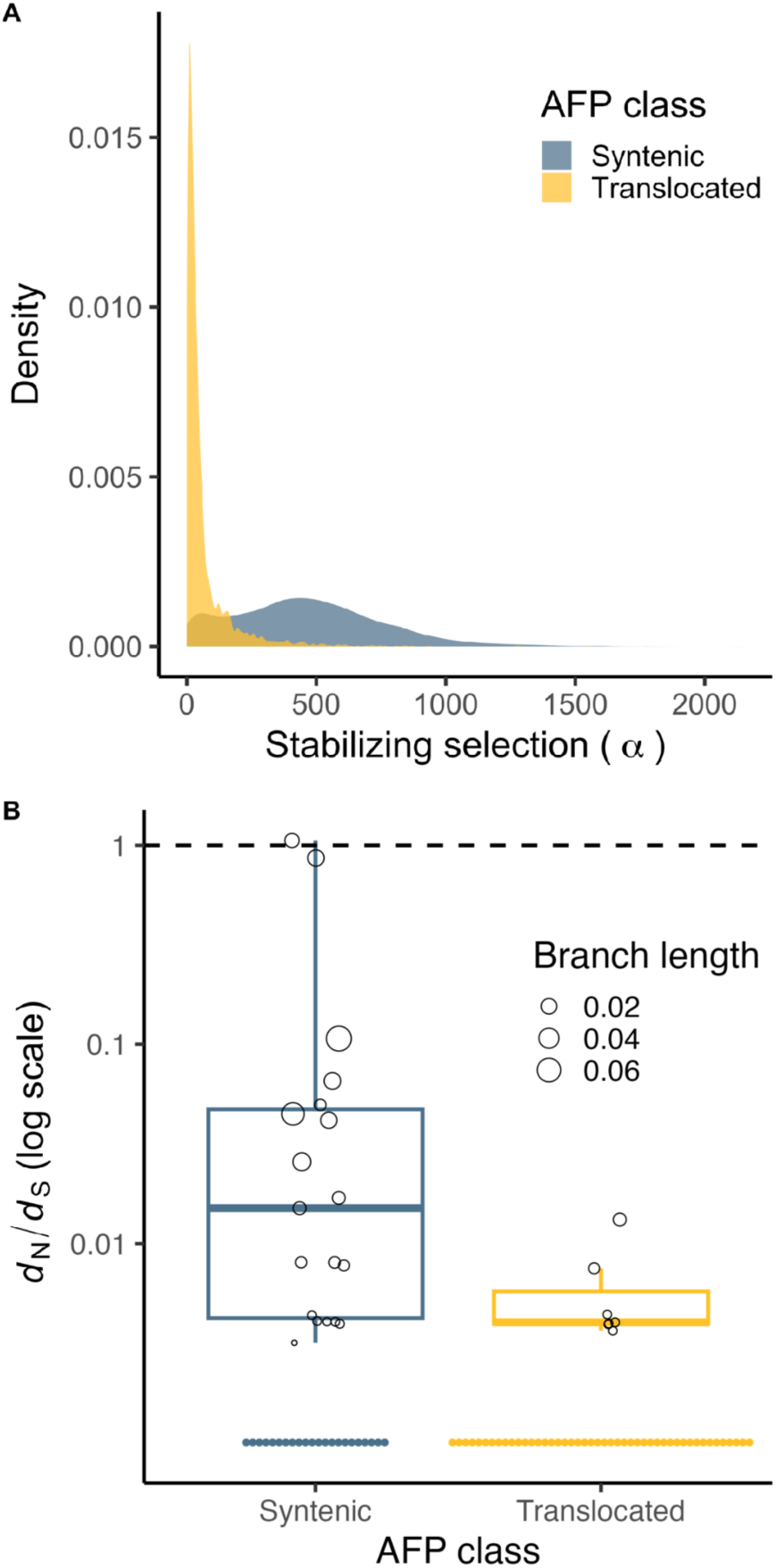
| Selection on type III antifreeze protein (AFP) copy number and coding sequence. (A) Posterior probabilities of stabilizing selection (**α**) on copy number of syntenic and translocated AFP III homologs across the Zoarcoidei tree derived from an Ornstein-Uhlenbeck phylogenetic mixed model. (B) Branch-tip *d*_N_/*d*_S_ ratios specific to syntenic and translocated AFP III homologs modeled in HyPhy RELAX. Each point represents a single gene. Branch length is depicted by point size. *d*_N_/*d*_S_ ratios of branch tips that lacked substitutions were undetermined and visualized below 0 as solid black circles.

Studies observing environmental effects on gene copy number are abundant (Kondrashov 2012; Lauer and Gresham 2019; Mérot et al. 2020), but the role of translocation in mediating environmental gradients of copy number is less understood. In populations of salmon, a chromosomal translocation shows strong genotype-environment association with temperature (Watson et al. 2022) while in soybean, a reciprocal chromosomal translocation may have facilitated climatic adaptation (Wang et al. 2021). Furthermore, translocation duplications of AFGP have been observed in notothenioid icefishes (Rivera-Colón et al. 2023), indicating potential for translocation to affect environmental gradients of antifreeze gene copy number in other systems. Our results expand on these studies by demonstrating that not only translocation, but its association with copy number, can contribute to molecular evolution across environments.

While our approach evaluated the hypothesis that AFPs expanded in shallow, polar zoarcoid species at a macroevolutionary scale, it is likely that environmental gradients in AFP copy number were attributed to intraspecific evolution as well. Population genomic studies of AFP III structural variation within Zoarcoidei species are necessary for further understanding environmental gradients in AFP copy number and the potential role of translocation in shaping these gradients. Intraspecific AFP copy number variation has been observed in populations of winter flounder inhabiting different depths and extents of sea ice cover (Hayes et al. 1991). Zoarcoids included in our study inhabit ranges that, in some cases, span both non-freezing and freezing conditions. Neutral and adaptive variation shaping AFP III copy number across the Zoarcoidei suborder can be more finely resolved by resequencing of conspecifics and populations distributed across gradients of temperature, pressure, and freezing risk. Detection and modeling of adaptive and neutral variation in AFP copy number is necessary for evaluating (i) additional evidence of whether copy number-environment associations arose via drift and (ii) whether non-neutral copy number variation is greater among translocated AFP III arrays.

## 3. Conclusion

Structural variation observable via long-read sequencing appears to have been critical for AFP III gene family evolution during the convergent adaptation of Zoarcoidei to both poles. Our findings highlight the role of multiple environmental stressors (temperature and pressure) in shaping the genetic architecture of adaptations like antifreeze function. Furthermore, structural variants such as translocations may act as gatekeepers of adaptation by constraining or facilitating molecular evolution under the selective pressures of those stressors. Future research should address whether translocation-associated gene family expansions are driven by mechanisms associated with transposable element architecture at translocation endpoints or break-induced replication. The reported, long-read Zoarcoidei reference genomes offer further potential to study the role of structural variation and chromosomal translocation in convergent environmental adaptation, as well as the complex evolution of antifreeze proteins.

## 4. Materials and Methods

### 4.1. DNA extraction, sequencing, and genome assembly

*Lycenchelys platyrhina* was collected from the Norwegian Sea, Ægir vent field, Norway (72.3286 °N, 1.5178 °E) on July 28th, 2021. The specimen for *Lycenchelys platyrhina* (originally described as *Lycodes platyrhinus* Jensen, 1902) received a putative identification during collection which required genomic identification and statistical consideration. *L. platyrhina* is a rarely observed species associated with only 11 occurrences in the Global Biodiversity Information Facility. The sequenced specimen was collected inside the range of *L. platyrhina* observations (67.1 – 73.8 °N, 10.3 °W – 8.2 °E). To this end, mitochondrial scaffolds of the *L. platyrhina* assembly were aligned to the NCBI nucleotide database using blastn (Altschul et al. 1990). No sequence of *Lycenchelys platyrhina* was available in the database. The top 10 greatest percent similarity blast hits were with species of the family Lycodinae to which *L. platyrhina* belongs; top 5 = *O. amberensis* (95.88%) *Lycenchelys kolthoffi* (95.57%), *Lycenchelys raridens* (95.05%), *Lycenchelys muraena* (94.60%), and *Lycenchelys sarsii* (94.60%). Skeletal muscle from a single individual was used for DNA isolation using the Circulomics Nanobind, BIG DNA prep, Tissue protocol. Isolated genomic DNA was quantified with a Fragment Analyzer (Agilent). Ampure-PB beads were used to purify and remove short fragments from the sample prior to library prep according to manufacturer protocol. A second size selection for >10 kb fragments was performed on the sample using a BluePippin system (Sage Science Inc., Beverly, MA, USA) with a 10kb cutoff. A library was prepared without shearing using SMRTbell® ExpressTemplate Prep Kit 3.0. Final library was size selected using BluePippin with an 8 kb cutoff. The library was sequenced on one 8M SMRT cells on Sequel IIe instrument using Sequel II Binding kit 2.2 and Sequencing Chemistry v2.0. Loading was performed by adaptive loading, movie time: 30 hours, pre-extension 2 hours.

Sampling of *Anarhichas lupus* was performed in July, 2022, on a captive-reared individual derived from eggs collected in Conception Bay, Newfoundland, Canada (47.697222 °N, – 52.921944 °W) via diving. Sampling of *Anarhichas minor* was performed in July, 2022, on a captive-reared individual bred from broodstock collected in St-Lawrence Gulf, Banc Beaugé, Canada (49.750000 °N, –60.166667 °W). Tissue was ground using dry ice. Cells were gently added to pre-warmed 0.8% agarose and mixture was placed into the wells of a pre-chilled plug mold. Following solidification, plugs were dislodged and treated with proteinase K and EDTA at 50°C. HMW DNA was extracted by dissolving the plugs at 70°C to remove the agarose followed by a phenol-chloroform extraction. Purification was performed on 15.0 μg of gDNA with AMPure PB beads and a bead-to-template ratio of 1. The DNA libraries were prepared following the Pacific Biosciences Preparing HiFi SMRTbell® Libraries using the SMRTbell Express Template Prep Kit 2.0 protocol. 10.0 μg of high molecular weight purified genomic DNA (final volume of 120 μl) was sheared using the Diagenode Megaruptor 3 instrument (Diagenode Inc., Denville, NJ, USA) and a two-cycle shear method at speed 33 followed by speed 34. The DNA Damage repair, End repair and SMRT bell ligation steps were performed as described in the template preparation protocol with the SMRTbell Template Prep Kit 2.0 reagents (Pacific Biosciences, Menlo Park, CA, USA). The DNA libraries were size selected on a BluePippin system using a 10 kb cutoff. The libraries underwent an AMPure PB bead cleanup following the SMRTlink v10 calculator procedure before being sequenced on a PacBio Sequel II instrument using adaptive loading, Sequel II Sequencing Kit 2.0, SMRT Cell 8M and 30 hours movies with a pre-extension time of 2h. Sequence data was processed using the SMRT Link Analysis v. 10.1.0.119588 pipeline.

*Bathymaster signatus* was collected in Alaska, USA (52.1861 °N, –173.596 °W), by the Alaska Trawl Survey on July 28th, 2012, via trawl net. DNA was extracted from gill filament via the Circulomics Nanobind Tissue Kit. HiFi Library prep was done with SMRTbell prep kit 3.0 (version 102-166-600 Apr 2022). As part of the library prep, the genomic DNA was sheared to 25kb on a Megaruptor 2.0. The library was size selected on the Blue Pippin according to PacBio’s recommended settings (10kb-50kb collection). The sample was sequenced on one SMRT cell on the Sequel II with v5 primer, v2.2 polymerase, 1 hr binding time, a 30 hr movie, 2 hr pre-extension, and adaptive loading.

*Lycodes diapterus* was collected in Monterey Bay, California, USA (36.8009 °N, – 121.903867 °W), on October, 20th, 2021. High molecular weight DNA was extracted from dissected muscle tissue using a Qiagen Genomic-tip DNA extraction kit. DNA was then sheared to ∼20 kbp using a Diagenode Megaruptor 3 and size selected for DNA for fragments greater than 10 kb using a BluePippin system. A PacBio HiFi library was prepared with the SMRTbell Express Template Prep Kit 2.0. The library was then sequenced on a PacBio Sequel II instrument on an 8M SMRT cell in CCS mode with a 30-hour movie time. DNA extraction, library preparation, and long-read sequencing of *Ophthalmolycus amberensis* are described in detail by Hotaling et al. (2023).

Haplotype-phased genomes of all species except for *O. amberensis* were assembled from long-read libraries using hifiasm v0.19.5-r592 (Cheng et al. 2021) set to default parameters. Assemblies of primary contigs are reported here. Read processing and genome assembly of *O. amberensis* was achieved using Canu (Koren et al. 2017) v1.8 and is described in detail in Hotaling, et al. (2023). Assembly completeness was evaluated using BUSCO v5.4.7 (Simão et al. 2015) input with universal, single-copy Actinopterygii orthologs. Collection coordinates for the specimens sequenced and reported in this study, and the *O. amberensis* specimen reported by Hotaling et al. (2023), are visualized in Figure S11.

### 4.2. AFP III and sasa/b annotation

Homologs of AFP III were annotated in each genome assembly included in the study using Exonerate v2.78.4 input with a query sequence of the translated *Macrozoarces americanus* AFP III (J03924.1) set to the model parameter ‘protein2genome’ (Slater and Birney 2005). *sasa*, *sasb*, and genes flanking the 5’ and 3’ ends of the ancestral syntenic region of Zoarcoidei AFP III (*sncgb* and *ldb3b*) were annotated using Exonerate (Slater and Birney 2005) set to the same parameters input with protein sequences of *sas* paralogs from *Lycodichthys dearborni* and *Gasterosteus aculeatus* protein sequences of *sncgb* (XP_040031911.1) and *ldb3b* (XP_040031907.1) as shown in Figure 3A. Exonerate BED file annotations of AFP III were filtered for whether hits contained start and stop codons and complete sequences of exons 1 and 2. Translocated AFP III copies outside of the ancestral syntenic AFP III region were manually reviewed for evidence of misassembly by aligning translocated AFP III regions to (i) known, characterized AFP translocation in *A. lupus* (Desjardins et al. 2012) and (ii) additional regions of translocation within single species if they possessed multiple translocation events to identify artificial duplications. This ancestral region is defined as the locus of conserved synteny at which AFP tandem arrays are flanked by the genes *sncgb* and *lbd3b*. Evidence of artificial AFP III translocations driven by misassembly was not apparent. Pseudogenized copies of AFP III were identified as regions mapping to AFP III query sequences that were complete but lacked a start codon, lacked one of the two AFP III exons, exhibited truncations of the majority of an exon, or conserved 5’/3’ untranslated regions that lacked a proximal AFP coding sequence. Publicly-available long-read genomes used in this study included *Cebidichthys violaceus* (GCA_023349555.1), *Gasterosteus aculeatus* (GCA_016920845.1), *Leptoclinus maculatus* (GCA_032191485.1), *Lycodes pacificus* (GCA_028022725.1), *Melanostigma gelatinosum* (GCA_949748355.1), *Pholis gunnellus* (GCA_910591455.2).

### 4.3. Environmental data

Depth-informed temperatures and salinities were mined from the Copernicus Global Ocean Reanalysis (GLORYS) database (Jean-Michel et al. 2021) within ±1° latitude/longitude of species’ coordinate of maximum likelihood of occurrence and the expected depth range of each species spanning a 10 year period between June 1st 2011 – June 1st 2021. Coordinates of maximum likelihood of occurrence were extracted from AquaMaps standardized range predictions (Kaschner et al. 2013), which are probabilistic predictions of abundance and occurrence derived from Global Biodiversity Information Facility occurrence data (Anon). AquaMaps ranges were constrained within coordinates of known species occurrences. Species minimum and maximum habitat depths were extracted from the FishBase (Froese 1996) depth range variable (Froese 1996). For *Lycenchelys platyrhina*, which lacked sufficient FishBase data, minimum and maximum observed depths were extracted from GBIF (Anon). Thermal minima were calculated as the minimum temperature experienced by each species within their representative latitude-longitude-depth range. Pressure was calculated as a function of sea water depth and latitude using conversions defined by the UNESCO Joint Panel on Oceanographic Tables and Standards (Fofonoff 1983) and expressed as a mean.

### 4.4. Species tree construction and phylogenetic modeling

A maximum likelihood species tree was constructed from MUSCLE v5 (Edgar 2004) alignments of concatenated protein sequences of 3,603 universal Actinopterygii single copy orthologs using IQ-Tree2 v1.6.12 parameterized with a WAG substitution model and fixed rate heterogeneity (Nguyen et al. 2015; Minh et al. 2020). A Bayesian phylogenetically-corrected model was then fitted to predict AFP III copy number variation as a function of species’ habitat temperature and pressure. The model assumed negative binomial variance for the AFP III count data and uninformative, uniform priors. Phylogenetic relatedness was fitted as a random effect of the genetic distance matrix derived from the species tree. The thermal minima and mean pressure of species’ habitats were included as fixed effects, with pressure fitted as a second order polynomial function. An iteration of this model was fitted that predicted AFP III copy number as a function of both pressure and depth. Because pressure and depth correlate, this model iteration was fitted to evaluate whether controlling for depth removed or reduced the effect of pressure. Pressure exhibited a significant and equivalent effect after controlling for depth, demonstrating that it is a stronger predictor of AFP copy number than depth. Model selection performed using Bayesian marginal likelihoods (Berger and Pericchi 1996) revealed that pressure was a better predictor of AFP III copy number than depth.

A multivariate iteration of the phylogenetic model was also fitted to predict two separate outcome variables for copy numbers of (i) AFP III within its ancestral, syntenic region and (ii) AFP III genes translocated out of the syntenic region. This multivariate model did not permit residual correlations. A third iteration of the phylogenetic model fit structural equations predicting variation in AFP III copy number as a function of phylogenetically independent variation in habitat thermal minimum and pressure. Phylogenetically independent variance in abiotic environments were estimated by predicting environmental variables as a function of phylogeny, and AFP III copy number as a function of environment and phylogeny. Thus, any effect of thermal minimum or pressure on AFP III copy number is quantified after conditioning on phylogenetic variation in abiotic environment.

Posterior sampling across all models was performed using four MCMC chains and 40,000 iterations including 10,000 warmups. All models were fitted using brms v2.20.4 (Bürkner 2017), an R interpreter of the Stan Bayesian programming language (Carpenter et al. 2017). The significance of fixed effects was assessed using a probability of direction test such that effects described as ‘significant’ exhibited posterior probabilities with 95% confidence intervals above or below 0 (Makowski, M.S. Ben-Shachar, et al. 2019). Because of potential misidentification of *L. platyrhina*, we evaluated the effect of the identification on model predictions. The leverage (the effect of a single observation on model predictions) that *L. platyrhina* imposed on parameter estimation in phylogenetic models of AFP gene family evolution across environments was quantified using a leave-one-out analysis. Removal of *L. platyrhina* from the univariate model imposed a 6.45% reduction in the effect of thermal minima on AFP III copy number and 4.24% reduction in the linear effect of pressure.

### 4.5. Analyses of AFP III gene family evolution and synteny

FASTA sequences of AFP III exons 1 and 2 and *sasa*/*b* exons 1 and 6 were extracted from annotation bed files using bed2fasta from bedtools v2.31.1 (Quinlan and Hall 2010). The two exons were concatenated per gene and input to a codon-aware alignment using PRANK v.170427 (Löytynoja 2014; Löytynoja 2021). A consensus gene tree was constructed from the alignment using IQ-TREE2 v1.6.12 set to fixed rate heterogeneity, 1000 bootstraps and optimization of a codon-to-protein substitution model, which selected a JTTDCMut+G4 model (Nguyen et al. 2015; Minh et al. 2020). Restricted maximum likelihood ancestral state reconstructions of AFP III copy number, habitat thermal minimum, and mean habitat pressure were performed using the ‘ace’ function of the R package ape v5.7-1 (Paradis et al. 2004; Paradis and Schliep 2019) in order to model contractions and expansions of AFP III across the studied phylogeny. A density tree visualizing all 1000 bootstrap gene trees and the consensus tree is plotted in Fig. S12. Ancestral state reconstructions of AFP III positions at the ancestral Zoarcoidei node were performed using ace(). Individual, discrete reconstructions were performed on 7 binary characters associated with whether species possessed putatively functional copies of AFP III at the syntenically-conserved locus or the six translocation loci. The relative likelihoods of the Zoarcoidei common ancestor possessing AFP III at these regions was measured as a ratio of maximum likelihoods for AFP III presence at the syntenically-conserved region divided by maximum likelihood of presence at each of the six translocation loci.

A multiple alignment of scaffolds or contigs containing translocated AFP III copies was performed using progressiveMAUVE v2.4.0 (Darling et al. 2010), which identified conservation and synteny of genomic regions conserved in at least two species of Zoarcoidei that exhibited translocated AFPs. This multiple alignment and the positions of translocated AFP arrays relative to conserved, syntenic regions are plotted in Fig. 3B in order to visualize conserved synteny of the ancestral AFP III locus, as well as the locations and number of AFP III translocations.

### 2.6. Models of selection on AFP III copy number and sequence

Stabilizing selection on AFP III copy number was modeled using Bayesian Ornstein-Uhlenbeck phylogenetic mixed models implemented in POUMM v.2.1.7 set to MCMC equals TRUE, 4 MCMC chains, and 80,000 samples (Mitov and Stadler 2018; Mitov and Stadler 2019). Significant differences between posterior probabilities of stabilizing selection on translocated and syntenic copy number were evaluated using a Bayes factor test implemented in bayestestR v0.13.2 (Makowski, M. Ben-Shachar, et al. 2019).

Purifying and diversifying selection were modeled across the AFP III gene tree by estimating branch-tip specific *d*_N_/*d*_S_ ratios in hyphy v2.5.62 RELAX (K. et al. 2005; Wertheim et al. 2015; Kosakovsky Pond et al. 2020). In this paper, we report branch-tip *d*_N_/*d*_S_ ratios as measures of purifying vs. diversifying selection, but we did not use RELAX to perform a statistical test of relaxed selection among a group of genes. As input to RELAX, we used a codon-aware alignment of all *sasa*, *sasb*, and AFP homologs produced using PRANK v.170427 (Löytynoja 2014; Löytynoja 2021) input with parameters –F and –codon and a guide tree produced using an IQ-Tree2 phylogenetic model fit to the PRANK alignment, parameterized with nucleotide substitution model optimization which selected a K2P+G4 model, fixed rate heterogeneity, and 1000 bootstraps (Edgar 2004; Löytynoja 2014; Löytynoja 2021). A nucleotide substitution model was selected because it produced more conservative *d*_N_/*d*_S_ estimates (i.e., less extreme absolute estimates of selection)(Katoh et al. 2002), which was used to produce the Figure 3C gene tree. Syntenic and translocated AFP homologs were defined as test and reference features with *sas* genes incorporated as background. Reported *d*_N_/*d*_S_ ratios were output by the RELAX general descriptive model (Wertheim et al. 2015). Variation in AFP *d*_N_/*d*_S_ was modeled as a function of whether copies fell within syntenic or translocated loci and gene branch tip lengths using gamma distributed Bayesian model specified as described under ‘Species tree construction and phylogenetic modeling’.

## Supporting information

Supplemental Results

## Acknowledgements

We acknowledge funding from National Science Foundation awards OPP-1543383 and OPP-1947040 (supporting TD), and OPP-2312253 (to JLK) as well as funding from the Antarctic Science Bursary. We acknowledge computing resources from the Center for Institutional Research Computing at Washington State University, the National Science Foundation Extreme Science and Engineering Discovery Environment (XSEDE), and University of California, Santa Cruz Research Computing. We thank the veterinary clinic staff of the Biodôme de Montréal, including Karine Béland, DMV. We also thank the captain and crew of the ARSV *Laurence M. Gould*, the personnel of the U.S. Antarctic Program Support Contractors for assistance in Chile, at sea and at Palmer Station, as well as the logistics in Denver, CO, for their support related to the Antarctic fieldwork required for this study. We acknowledge funding from the Trond Mohn Science Foundation through the Centre for Deep Sea Research (grant number TMS2020TMT13) and the Norwegian Biodiversity Information Centre (the Taxonomy Initiative, project nr. 3-20-70184243) and thank the scientists and students from University of Bergen (Norway), the crew of the RV GO Sars and the pilots of the ROV ÆGIR6000 for their assistance with sampling in the Arctic. Lastly, we would like to thank the Alaskan Trawl Survey, and the Ocean Genome Legacy Center, Northeastern University, for assisting with specimen collections and archiving.

## Author Contributions

Designed research (JLK, SNB)

Performed research (FGH, LSFL, IB, JLK, MHE, NRLF, NS, PBF, SH, SNB, TA, TD)

Analyzed data (JLK, NS, SNB) Wrote the paper (JLK, NS, SNB)

Edited the paper (FGH, LSFL, IB, JLK, MHE, NRLF, NS, PBF, SH, SNB, TA, TD)

## Data Availability

Genome assemblies, raw sequencing reads, and associated files are available on NCBI under the following Bioprojects: PRJNA980960 (*Anarhichas lupus*), PRJNA982125 (*Anarhichas minor*), PRJNA982427 (*Bathymaster signatus*), PRJNA985917 (*Lycenchelys platyrhina*), PRJNA982746 (*Lycodes diapterus*) and PRJNA701078 (*Ophalmolycus amberensis*). All code used for bioinformatic processing and statistical analysis are available under the GitHub repository https://github.com/snbogan/Long_AFP [available to reviewers upon request or acceptance and publication] in addition to intermediate data files and specimen metadata. An archived copy of this repository is available on Zenodo under the DOI [to be generated upon acceptance].

## References

1. Altschul SF, Gish W, Miller W, Myers EW, Lipman DJ. 1990. Basic local alignment search tool. J. Mol. Biol. 215:403–410.

2. Anderson ME. 1994. Systematics and osteology of the Zoarcidae (Teleostei: Perciformes). *JLB Smith Institute of Ichthyology*, Ichthyological Bulletin 60:1–120.

3. Anon. GBIF.org. Global Biodiversity Information Facility [Internet]. Available from: https://www.gbif.org

4. Auger M, Morrow R, Kestenare E, Sallée J-B, Cowley R. 2021. Southern Ocean in-situ temperature trends over 25 years emerge from interannual variability. Nat. Commun. 12:514.

5. Baalsrud HT, Tørresen OK, Solbakken MH, Salzburger W, Hanel R, Jakobsen KS, Jentoft S. 2018. De Novo Gene Evolution of Antifreeze Glycoproteins in Codfishes Revealed by Whole Genome Sequence Data. Mol. Biol. Evol. 35:593–606.

6. Barrett J. 2001. Thermal hysteresis proteins. Int. J. Biochem. Cell Biol. 33:105–117.

7. Baskaran A, Kaari M, Venugopal G, Manikkam R, Joseph J, Bhaskar PV. 2021. Anti freeze proteins (Afp): Properties, sources and applications – A review. Int. J. Biol. Macromol. 189:292–305.

8. Berger JO, Pericchi LR. 1996. The Intrinsic Bayes Factor for Model Selection and Prediction. J. Am. Stat. Assoc. 91:109–122.

9. Bista I, Collins M, Wellcome Sanger Institute Tree of Life Management, Samples and Laboratory team. 2024. The genome sequence of the limp eelpout, Melanostigma gelatinosum Günther, 1881. Wellcome Open Research 9:192.

10. Bista I, Wood JMD, Desvignes T, McCarthy SA, Matschiner M, Ning Z, Tracey A, Torrance J, Sims Y, Chow W, et al. 2023. Genomics of cold adaptations in the Antarctic notothenioid fish radiation. Nat. Commun. 14:3412.

11. Bodnar RJ. 1993. Revised equation and table for determining the freezing point depression of H2O-Nacl solutions. Geochim. Cosmochim. Acta 57:683–684.

12. Bürkner P-C. 2017. brms: An R Package for Bayesian Multilevel Models Using Stan. J. Stat. Softw. 80:1–28.

13. Carpenter B, Gelman A, Hoffman MD, Lee D, Goodrich B, Betancourt M, Brubaker MA, Guo J, Li P, Riddell A. 2017. Stan: A Probabilistic Programming Language. J. Stat. Softw. [Internet] 76. Available from: 10.18637/jss.v076.i01

14. Celik Y, Graham LA, Mok Y-F, Bar M, Davies PL, Braslavsky I. 2010. Superheating of ice crystals in antifreeze protein solutions. Proc. Natl. Acad. Sci. U. S. A. 107:5423–5428.

15. Cheng C, Devries A. 2002. Origins and evolution of fish antifreeze proteins. Fish antifreeze proteins:83–107.

16. Cheng H, Concepcion GT, Feng X, Zhang H, Li H. 2021. Haplotype-resolved de novo assembly using phased assembly graphs with hifiasm. Nat. Methods 18:170–175.

17. Cziko PA, DeVries AL, Evans CW, Cheng C-HC. 2014. Antifreeze protein-induced superheating of ice inside Antarctic notothenioid fishes inhibits melting during summer warming. Proc. Natl. Acad. Sci. U. S. A. 111:14583–14588.

18. Darling AE, Mau B, Perna NT. 2010. progressiveMauve: multiple genome alignment with gene gain, loss and rearrangement. PLoS One 5:e11147.

19. Davies PL. 2022. Reflections on antifreeze proteins and their evolution. Biochem. Cell Biol. 100:282–291.

20. Deng C, Cheng C-HC, Ye H, He X, Chen L. 2010. Evolution of an antifreeze protein by neofunctionalization under escape from adaptive conflict. Proc. Natl. Acad. Sci. U. S. A. 107:21593–21598.

21. Desjardins M, Graham LA, Davies PL, Fletcher GL. 2012. Antifreeze protein gene amplification facilitated niche exploitation and speciation in wolffish. FEBS J. 279:2215–2230.

22. DeVries AL. 1971. Glycoproteins as biological antifreeze agents in antarctic fishes. Science 172:1152–1155.

23. DeVries AL, Cheng C-H C. 2005. Antifreeze Proteins and Organismal Freezing Avoidance in Polar Fishes. In: Fish Physiology. Vol. 22. Academic Press. p. 155–201.

24. DeVries AL, Wohlschlag DE. 1969. Freezing resistance in some Antarctic fishes. Science 163:1073–1075.

25. Duman JG. 2015. Animal ice-binding (antifreeze) proteins and glycolipids: an overview with emphasis on physiological function. J. Exp. Biol. 218:1846–1855.

26. Edgar RC. 2004. MUSCLE: multiple sequence alignment with high accuracy and high throughput. Nucleic Acids Res. 32:1792–1797.

27. Fetterer F;. KK;. MW;. SM;. WA. 2017. Sea Ice Index, Version 3. Available from: https://nsidc.org/data/g02135/versions/3

28. Fofonoff NP. 1983. Algorithms for the computation of fundamental properties of seawater. Available from: https://repository.oceanbestpractices.org/handle/11329/109

29. Friedman ST, Muñoz MM. 2023. A latitudinal gradient of deep-sea invasions for marine fishes. Nat. Commun. 14:773.

30. Froese R. 1996. FishBase. Oceanographic Literature Review 3:321.

31. Fujino K, Lewis EL, Perkin RG. 1974. The freezing point of seawater at pressures up to 100 bars. J. Geophys. Res. 79:1792–1797.

32. Gilbert JA, Davies PL, Laybourn-Parry J. 2005. A hyperactive, Ca2+-dependent antifreeze protein in an Antarctic bacterium. FEMS Microbiol. Lett. 245:67–72.

33. Gillett MB, Suko JR, Santoso FO, Yancey PH. 1997. Elevated levels of trimethylamine oxide in muscles of deep-sea gadiform teleosts: A high-pressure adaptation? J. Exp. Zool. 279:386– 391.

34. Graham LA, Davies PL. 2024. Fish antifreeze protein origin in sculpins by frameshifting within a duplicated housekeeping gene. FEBS J. 291:4043–4061.

35. Griffith RW. 1981. Composition of the blood serum of deep-sea fishes. Biol. Bull. 160:250–264.

36. Gupta R, Deswal R. 2014. Antifreeze proteins enable plants to survive in freezing conditions. J. Biosci. 39:931–944.

37. Harewood L, Schütz F, Boyle S, Perry P, Delorenzi M, Bickmore WA, Reymond A. 2010. The effect of translocation-induced nuclear reorganization on gene expression. Genome Res. 20:554–564.

38. Hargens AR. 1972. Freezing resistance in polar fishes. Science 176:184–186.

39. Hastings PJ, Ira G, Lupski JR. 2009. A microhomology-mediated break-induced replication model for the origin of human copy number variation. PLoS Genet. 5:e1000327.

40. Hastings PJ, Lupski JR, Rosenberg SM, Ira G. 2009. Mechanisms of change in gene copy number. Nat. Rev. Genet. 10:551–564.

41. Hayes PH, Davies PL, Fletcher GL. 1991. Population differences in antifreeze protein gene copy number and arrangement in winter flounder. Genome 34:174–177.

42. Hew CL, Wang NC, Joshi S, Fletcher GL, Scott GK, Hayes PH, Buettner B, Davies PL. 1988. Multiple genes provide the basis for antifreeze protein diversity and dosage in the ocean pout, Macrozoarces americanus. J. Biol. Chem. 263:12049–12055.

43. Hickerson MJ, Cunningham CW. 2006. Nearshore fish (*Pholis gunnellus*) persists across the North Atlantic through multiple glacial episodes: N. ATLANTIC PERSISTENCE INPHOLIS GUNNELLUS. Mol. Ecol. 15:4095–4107.

44. Hobbs RS, Hall JR, Graham LA, Davies PL, Fletcher GL. 2020. Antifreeze protein dispersion in eelpouts and related fishes reveals migration and climate alteration within the last 20 Ma. PLoS One 15:e0243273.

45. Hotaling S, Borowiec ML, Lins LSF, Desvignes T, Kelley JL. 2021. The biogeographic history of eelpouts and related fishes: Linking phylogeny, environmental change, and patterns of dispersal in a globally distributed fish group. Mol. Phylogenet. Evol. 162:107211.

46. Hotaling S, Desvignes T, Sproul JS, Lins LSF, Kelley JL. 2023. Pathways to polar adaptation in fishes revealed by long-read sequencing. Molecular Ecology 32:1381–1397.

47. Hull RM, Cruz C, Jack CV, Houseley J. 2017. Environmental change drives accelerated adaptation through stimulated copy number variation. PLoS Biol. 15:e2001333.

48. Jean-Michel L, Eric G, Romain B-B, Gilles G, Angélique M, Marie D, Clément B, Mathieu H, Olivier LG, Charly R, et al. 2021. The Copernicus Global 1/12° Oceanic and Sea Ice GLORYS12 Reanalysis. Front Earth Sci. Chin. [Internet] 9. Available from: https://www.frontiersin.org/articles/10.3389/feart.2021.698876

49. Jia Z, DeLuca CI, Chao H, Davies PL. 1996. Structural basis for the binding of a globular antifreeze protein to ice. Nature 384:285–288.

50. Kaschner K, Rius-Barile J, Kesner-Reyes K. 2013. About AquaMaps. Available from: https://www.aquamaps.org/AboutAquaMaps.htm

51. Katoh K, Misawa K, Kuma K-I, Miyata T. 2002. MAFFT: a novel method for rapid multiple sequence alignment based on fast Fourier transform. Nucleic Acids Res. 30:3059–3066.

52. Kelley JL, Aagaard JE, MacCoss MJ, Swanson WJ. 2010. Functional diversification and evolution of antifreeze proteins in the antarctic fish Lycodichthys dearborni. J. Mol. Evol. 71:111–118.

53. Kidwell MG. 2002. Transposable elements and the evolution of genome size in eukaryotes. Genetica 115:49–63.

54. Kondrashov FA. 2012. Gene duplication as a mechanism of genomic adaptation to a changing environment. Proc. Biol. Sci. 279:5048–5057.

55. Koren S, Walenz BP, Berlin K, Miller JR, Bergman NH, Phillippy AM. 2017. Canu: scalable and accurate long-read assembly via adaptive k-mer weighting and repeat separation. Genome Res. 27:722–736.

56. Kosakovsky Pond SL, Poon AFY, Velazquez R, Weaver S, Hepler NL, Murrell B, Shank SD, Magalis BR, Bouvier D, Nekrutenko A, et al. 2020. HyPhy 2.5-A customizable platform for evolutionary hypothesis testing using PHYlogenies. Mol. Biol. Evol. 37:295–299.

57. K. PSL, Frost SD, Muse SV. 2005. HyPhy: hypothesis testing using phylogenies. Bioinformatics 21:676–679.

58. Kramara J, Osia B, Malkova A. 2018. Break-Induced Replication: The Where, The Why, and The How. Trends Genet. 34:518–531.

59. Lauer S, Gresham D. 2019. An evolving view of copy number variants. Curr. Genet. 65:1287– 1295.

60. Löytynoja A. 2014. Phylogeny-aware alignment with PRANK. Methods Mol. Biol. 1079:155–170.

61. Löytynoja A. 2021. Phylogeny-aware alignment with PRANK and PAGAN. Methods Mol. Biol. 2231:17–37.

62. Makowski D, Ben-Shachar M, Lüdecke D. 2019. BayestestR: Describing effects and their uncertainty, existence and significance within the Bayesian framework. J. Open Source Softw. 4:1541.

63. Makowski D, Ben-Shachar MS, Chen SHA, Lüdecke D. 2019. Indices of Effect Existence and Significance in the Bayesian Framework. Front. Psychol. 10:2767.

64. Marx V. 2023. Method of the year: long-read sequencing. Nat. Methods 20:6–11.

65. Mérot C, Oomen RA, Tigano A, Wellenreuther M. 2020. A roadmap for understanding the evolutionary significance of structural genomic variation. Trends Ecol. Evol. 35:561–572.

66. Minh BQ, Schmidt HA, Chernomor O, Schrempf D, Woodhams MD, von Haeseler A, Lanfear R. 2020. IQ-TREE 2: New Models and Efficient Methods for Phylogenetic Inference in the Genomic Era. Mol. Biol. Evol. 37:1530–1534.

67. Mitov V, Stadler T. 2018. A practical guide to estimating the heritability of pathogen traits. Mol. Biol. Evol. 35:756–772.

68. Mitov V, Stadler T. 2019. Parallel likelihood calculation for phylogenetic comparative models: The SPLITT C++ library. Methods Ecol. Evol. 10:493–506.

69. Møller PR, Nielsen JG, Anderson ME. 2005. Systematics of Polar Fishes. In: Fish Physiology. Vol. 22. Academic Press. p. 25–78.

70. Nath S, Shaw DE, White MA. 2021. Improved contiguity of the threespine stickleback genome using long-read sequencing. G3 (Bethesda) 11:jkab007.

71. Nguyen L-T, Schmidt HA, von Haeseler A, Minh BQ. 2015. IQ-TREE: a fast and effective stochastic algorithm for estimating maximum-likelihood phylogenies. Mol. Biol. Evol. 32:268–274.

72. Paradis E, Claude J, Strimmer K. 2004. APE: Analyses of Phylogenetics and Evolution in R language. Bioinformatics 20:289–290.

73. Paradis E, Schliep K. 2019. ape 5.0: an environment for modern phylogenetics and evolutionary analyses in R. Bioinformatics 35:526–528.

74. Ponte RM. 2013. Heat content and temperature of the ocean. In: Earth System Monitoring. New York, NY: Springer New York. p. 153–180.

75. Potter RW, Clynne MA, Brown DL. 1978. Freezing point depression of aqueous sodium chloride solutions. Econ. Geol. 73:284–285.

76. Programme WSIT of L, Potter S, Collective DT of LB, Collective WSISODP, Collective T of LCI, of Life Consortium DT. 2022. The genome sequence of the rock gunnel, Pholis gunnellus (Linnaeus, 1758). Wellcome Open Res. 7:58.

77. Quinlan AR, Hall IM. 2010. BEDTools: a flexible suite of utilities for comparing genomic features. Bioinformatics 26:841–842.

78. Radchenko OA. 2016. Timeline of the evolution of eelpouts from the suborder Zoarcoidei (Perciformes) based on DNA variability. Journal of Ichthyology 56:556–568.

79. Raymond JA, DeVries AL. 1977. Adsorption inhibition as a mechanism of freezing resistance in polar fishes. Proc. Natl. Acad. Sci. U. S. A. 74:2589–2593.

80. Reist JD, Wrona FJ, Prowse TD, Power M, Dempson JB, Beamish RJ, King JR, Carmichael TJ, Sawatzky CD. 2006. General effects of climate change on Arctic fishes and fish populations. Ambio 35:370–380.

81. Rhie A, McCarthy SA, Fedrigo O, Damas J, Formenti G, Koren S, Uliano-Silva M, Chow W, Fungtammasan A, Kim J, et al. 2021. Towards complete and error-free genome assemblies of all vertebrate species. Nature 592:737–746.

82. Rivera-Colón AG, Rayamajhi N, Minhas BF, Madrigal G, Bilyk KT, Yoon V, Hüne M, Gregory S, Cheng CHC, Catchen JM. 2023. Genomics of secondarily temperate adaptation in the only non-Antarctic icefish. Mol. Biol. Evol. 40:msad029.

83. Rives N, Lamba V, Cheng CHC, Zhuang X. 2024. Diverse origins of near-identical antifreeze proteins in unrelated fish lineages provide insights into evolutionary mechanisms of new gene birth and protein sequence convergence. Mol. Biol. Evol. 41:msae182.

84. Schmidt JM, Good RT, Appleton B, Sherrard J, Raymant GC, Bogwitz MR, Martin J, Daborn PJ, Goddard ME, Batterham P, et al. 2010. Copy number variation and transposable elements feature in recent, ongoing adaptation at the Cyp6g1 locus. PLoS Genet. 6:e1000998.

85. Scholander PF, Flagg W, Hock RJ, Irving L. 1953. Studies on the physiology of frozen plants and animals in the arctic. J. Cell. Comp. Physiol. 42:1–56.

86. Simão FA, Waterhouse RM, Ioannidis P, Kriventseva EV, Zdobnov EM. 2015. BUSCO: assessing genome assembly and annotation completeness with single-copy orthologs. Bioinformatics 31:3210–3212.

87. Slater GSC, Birney E. 2005. Automated generation of heuristics for biological sequence comparison. BMC Bioinformatics 6:31.

88. Storey KB, Storey JM. 1988. Freeze tolerance in animals. Physiol. Rev. 68:27–84.

89. Verde C, Parisi E, di Prisco G. 2006. The evolution of thermal adaptation in polar fish. Gene 385:137–145.

90. Wang W, Chen L, Wang X, Duan J, Flynn RD, Wang Y, Clark CB, Sun L, Zhang D, Wang DR, et al. 2021. A transposon-mediated reciprocal translocation promotes environmental adaptation but compromises domesticability of wild soybeans. New Phytol. 232:1765–1777.

91. Wang Y, Wang X, Paterson AH. 2012. Genome and gene duplications and gene expression divergence: a view from plants. Ann. N. Y. Acad. Sci. 1256:1–14.

92. Watson KB, Lehnert SJ, Bentzen P, Kess T, Einfeldt A, Duffy S, Perriman B, Lien S, Kent M, Bradbury IR. 2022. Environmentally associated chromosomal structural variation influences fine-scale population structure of Atlantic Salmon (Salmo salar). Mol. Ecol. 31:1057–1075.

93. Watson SK, deLeeuw RJ, Horsman DE, Squire JA, Lam WL. 2007. Cytogenetically balanced translocations are associated with focal copy number alterations. Hum. Genet. 120:795–805.

94. Wertheim JO, Murrell B, Smith MD, Kosakovsky Pond SL, Scheffler K. 2015. RELAX: detecting relaxed selection in a phylogenetic framework. Mol. Biol. Evol. 32:820–832.

95. Worster MG, Rees Jones DW. 2015. Sea-ice thermodynamics and brine drainage. Philos. Trans. A Math. Phys. Eng. Sci. 373:20140166.

96. Wright DB, Escalona M, Marimuthu MPA, Sahasrabudhe R, Nguyen O, Sacco S, Beraut E, Toffelmier E, Miller C, Shaffer HB, et al. 2023. Reference genome of the Monkeyface Prickleback, Cebidichthys violaceus. J. Hered. 114:52–59.

97. Yancey PH, Blake WR, Conley J. 2002. Unusual organic osmolytes in deep-sea animals: adaptations to hydrostatic pressure and other perturbants. Comp. Biochem. Physiol. A Mol. Integr. Physiol. 133:667–676.

98. Yancey PH, Gerringer ME, Drazen JC, Rowden AA, Jamieson A. 2014. Marine fish may be biochemically constrained from inhabiting the deepest ocean depths. Proc. Natl. Acad. Sci. U. S. A. 111:4461–4465.

99. Zhuang X, Yang C, Murphy KR, Cheng C-HC. 2019. Molecular mechanism and history of non-sense to sense evolution of antifreeze glycoprotein gene in northern gadids. Proc. Natl. Acad. Sci. U. S. A. 116:4400–4405.

