## Supplemental Results for "Temperature and Pressure Shaped the Evolution of Antifreeze Proteins in Polar and Deep Sea Zoarcoid Fishes"

| Species | Syntenic AFP copies | Translocated AFP copies | Habitat thermal minima (°C) | Meann habitat pressure (bars) |
|---|---|---|---|---|
| *Anarhichas lupus* | 5 | 13 | 2.94744706 | 34.3407065 |
| *Anarhichas minor* | 3 | 2 | 3.50044251 | 35.1212676 |
| *Bathymaster signatus* | 0 | 0 | -1.8280892 | 33.9869844 |
| *Cebidichthys violaceus* | 0 | 0 | 7.20880747 | 31.6912469 |
| *Lycodes diapterus* | 0 | 0 | -0.4921109 | 34.035235 |
| *Leptoclinus maculatus* | 4 | 38 | -1.9408857 | 33.9058768 |
| *Lycodes pacificus* | 0 | 0 | 5.10156584 | 33.4709885 |
| *Lycenchelys platyrhina* | 1 | 0 | -1.0026245 | 34.9135639 |
| *Melanostigma gelatinosum* | 0 | 0 | -1.5695364 | 34.2924081 |
| *Ophthalmolycus amberensis* | 13 | 0 | -2.9992676 | 34.5588572 |
| *Pholis gunnellus* | 15 | 0 | 2.51017785 | 34.6078114 |

**Table S1** | Zoarcoidei species names, antifreeze protein III copy numbers, habitat thermal minima, and mean habitat pressures.

| Species | Family | Complete AFP III sequences | Partial AFP III sequences | NCBI Genome accession |
|---|---|---|---|---|
| *Anoplopoma fimbria* | Anoplopomatidae | 0 | 0 | GCA_027596085.2 |
| *Clinocottus analis* | Cottidae | 0 | 0 | GCA_023055335.1 |
| *Cottus gobio* | Cottidae | 0 | 0 | GCA_023566465.1 |
| *Cottus rhenanus* | Cottidae | 0 | 0 | GCA_001455555.1 |
| *Myoxocephalus scorpius* | Cottidae | 0 | 0 | GCA_900312955.1 |
| *Taurulus bubalis* | Cottidae | 0 | 0 | GCA_021346845.1 |
| *Cyclopterus lumpus* | Cyclopteridae | 0 | 0 | GCA_009769545.1 |
| *Apeltes quadracus* | Gasterosteidae | 0 | 0 | GCA_021346845.1 |
| *Ophiodon elongatus* | Hexagrammidae | 0 | 0 | GCA_016806645.1 |
| *Liparis tanakae* | Liparidae | 0 | 0 | GCA_036178185.1 |
| *Pseudoliparis sp. Yap Trench* | Liparidae | 0 | 0 | GCA_004335475.1 |
| *Pseudoliparis swirei* | Liparidae | 0 | 0 | GCA_029220125.1 |

**Table S2** | Absence of complete and partial type III antifreeze protein (AFP III) genes in non-zoarcoid members of Scorpaeniformes. AFP III sequences were screened for and annotated using the same methods employed for Zoarcoidei and *Gasterosteus aculeatus*.

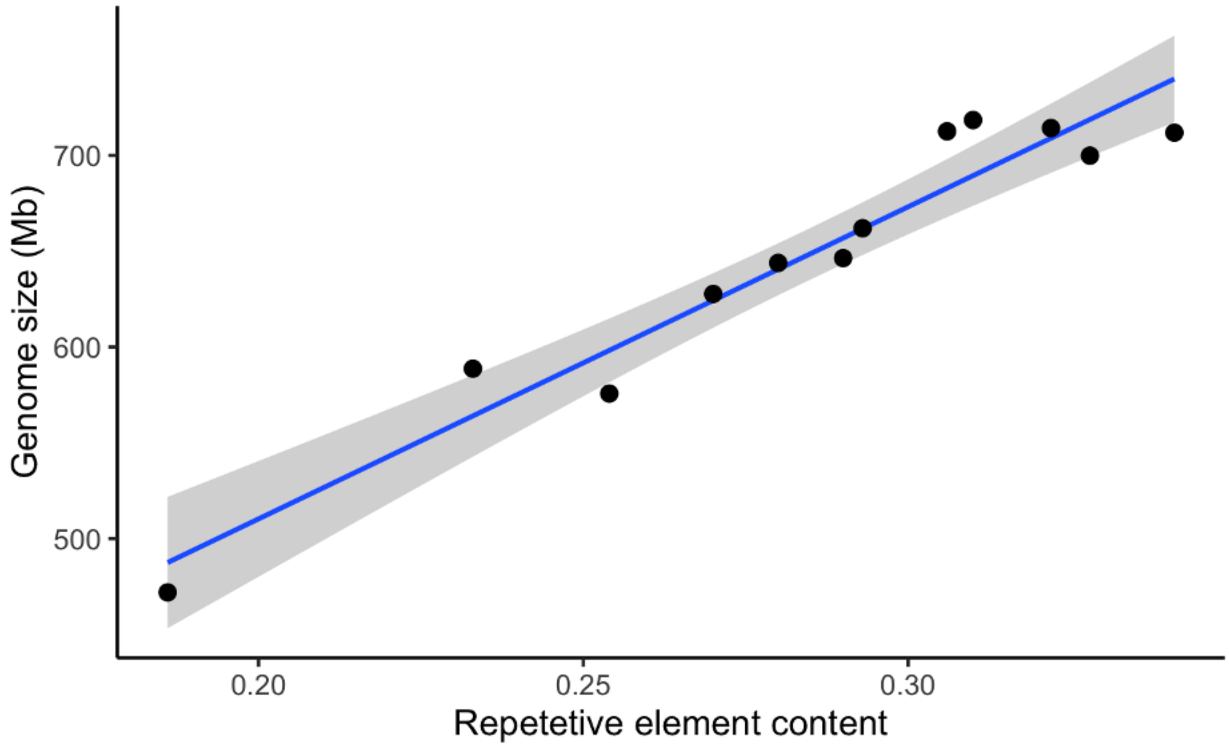


**Figure S1** | Correlation between genome size and proportions of repetitive element content in Zoarcoidei and *Gasterosteus aculeatus* outgroup. Points represent all zoarcoid species included in this study ($r^2 = 0.93$).

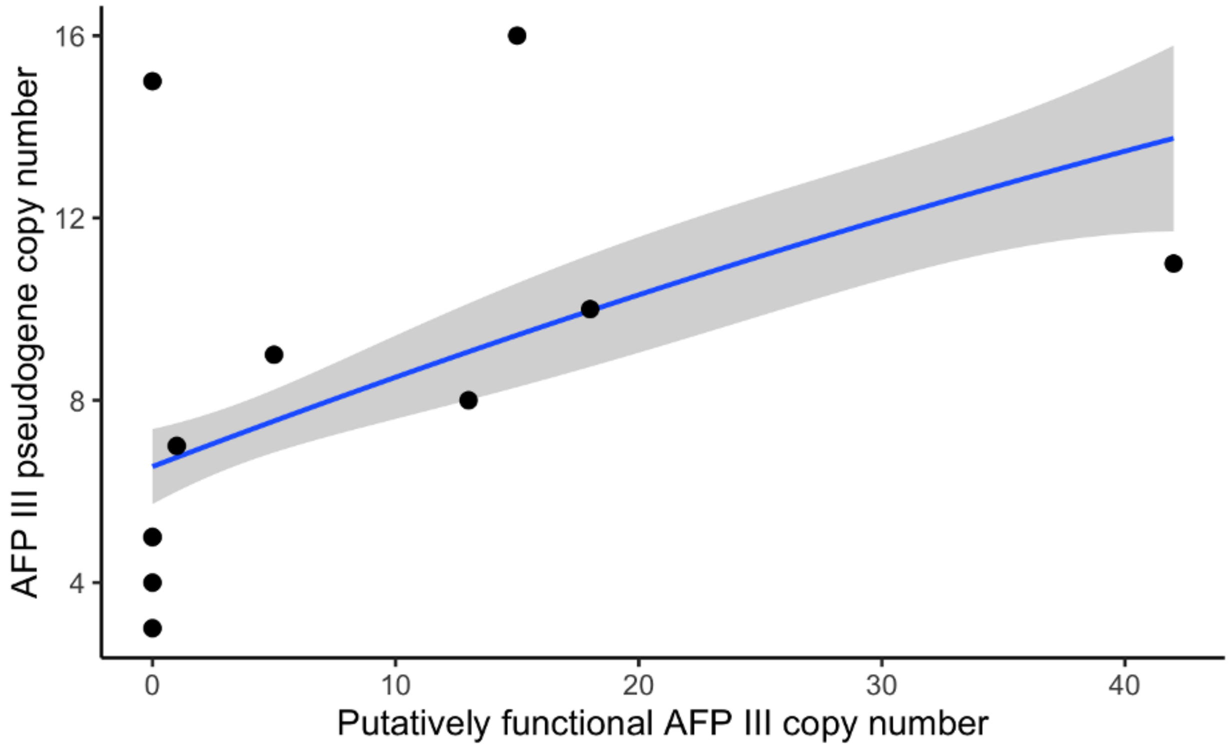


**Figure S2** | Non-linear correlation between putatively functional and pseudogenized copies of Zoarcoidei type III antifreeze proteins ($r^2 = 0.31$). Points represent all zoarcoid species included in this study. Also plotted is a best fitted curve of a second-order polynomial derived from a phylogenetically-controlled model.

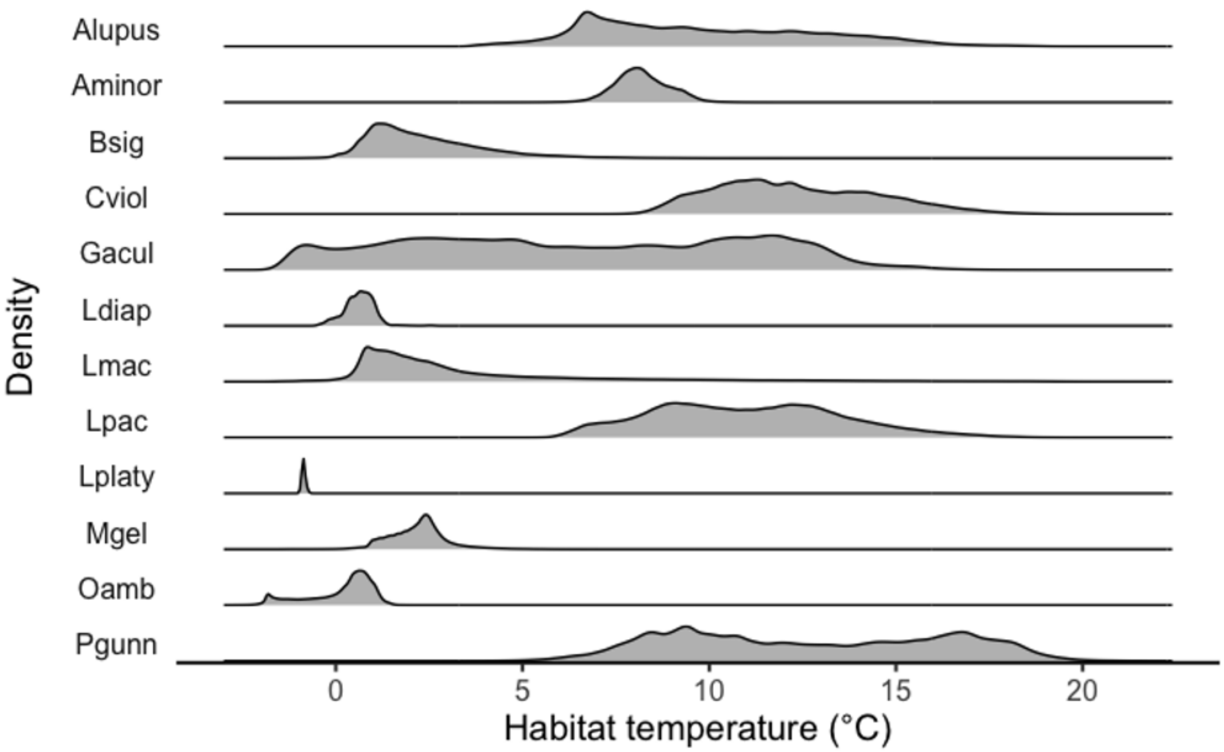


**Figure S3** | Distributions of species' depth-informed habitat temperatures extracted from the Copernicus Global Ocean Reanalysis Survey: June 1st, 2011 – June 1st, 2021. Y axes are labeled with abbreviated species names (Alupus = *Anarhichas lupus*; Aminor = *Anarhichas minor*; Bsig = *Bathymaster signatus*; Cviol = Cebidichthys violaceus; Gacul = *Gasterosteus aculeatus*; Ldiap = *Lycodes diapterus*; Lmac = *Leptoclinus maculatus*; Lpac = *Lycodes pacificus*; Lplaty = *Lycenchelys platyrhina*; Mgel = *Melanostigma gelatinosum*; Oamb = *Opthalmolycus amberensis*; Pgunn = *Pholis gunnellus*).

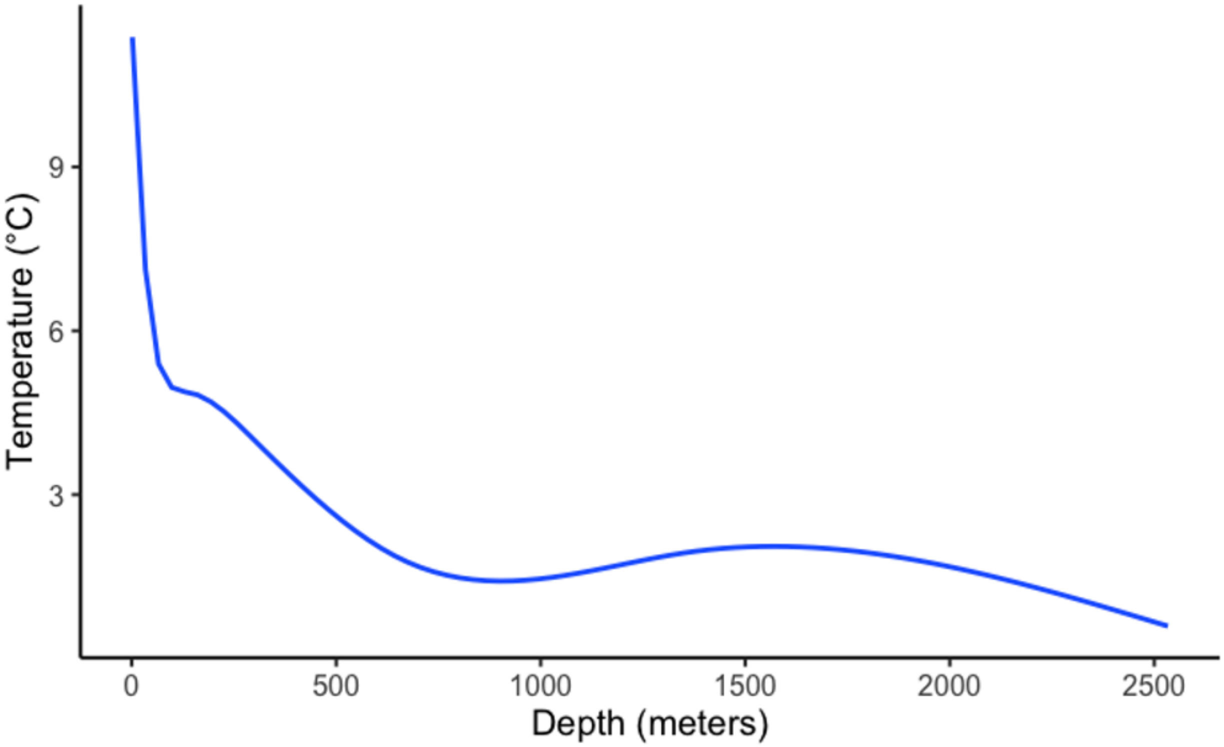


**Figure S4** | Correlation between mean temperature and depth in Copernicus Global Ocean Reanalysis (GLORYs) dataset. A nonlinear, loess curve is plotted that was fitted to GLORYs temperature and depth data from Zoarcoidei habitats.

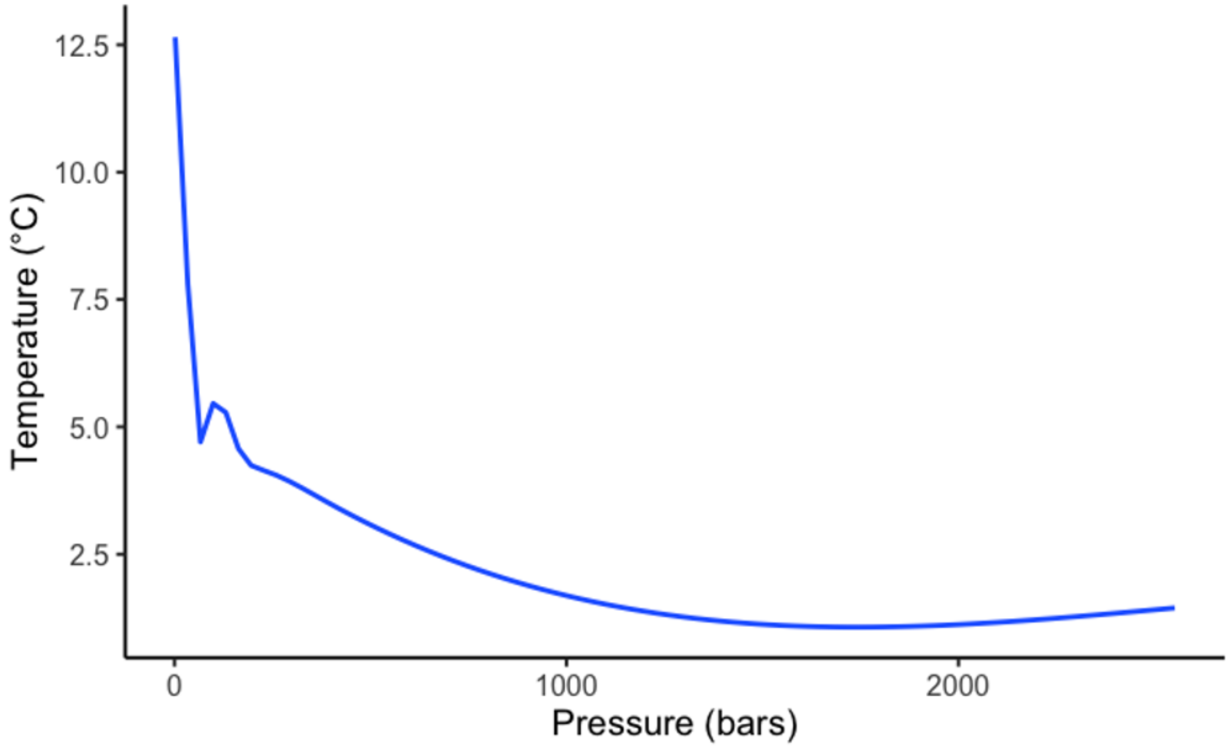


**Figure S5** | Correlation between mean temperature and pressure, calculated from Copernicus Global Ocean Reanalysis (GLORYs) data. A nonlinear, loess curve is plotted that was fitted to GLORYs temperatures and inferred pressure across Zoarcoidei habitats.

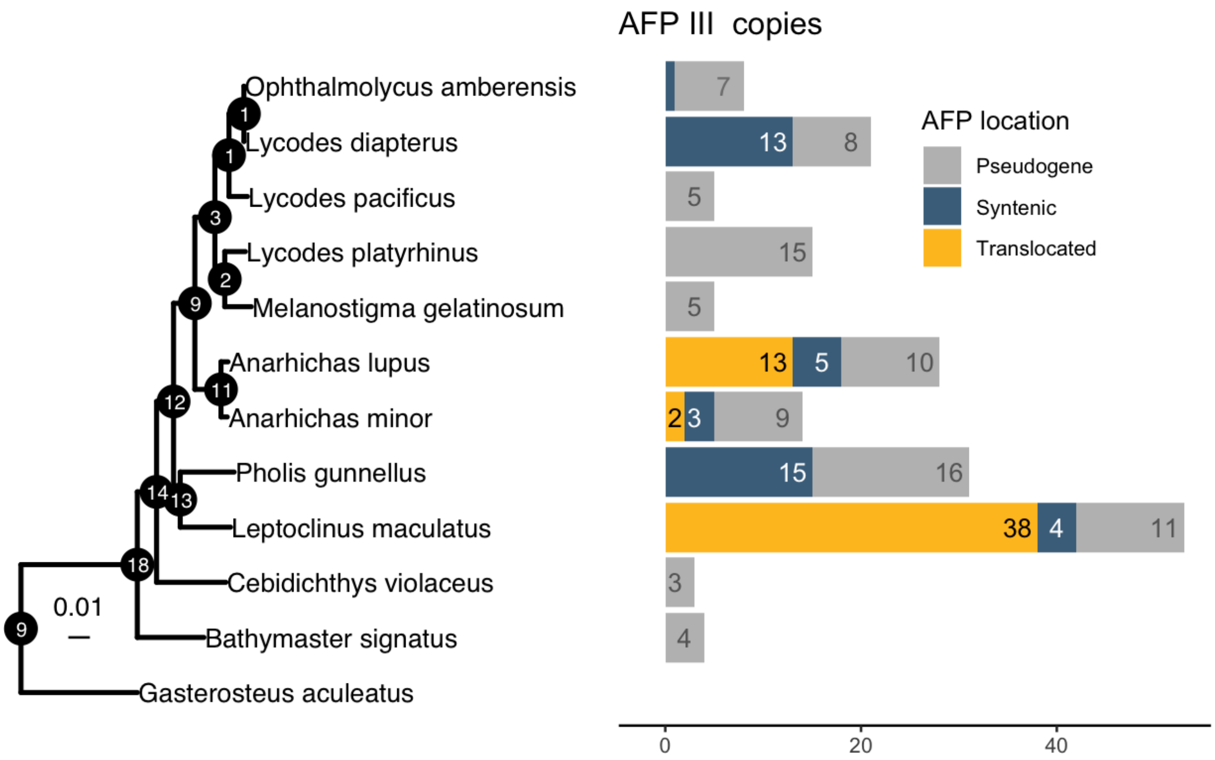


**Figure S6** | Ancestral state reconstruction of putatively functional (non-pseudogenized) type III antifreeze protein (AFP III) copy number. Predicted ancestral copy number is enumerated at tree nodes. Branch lengths represent substitutions per base pair. Copies of AFP III present in the ancestral syntenic region, present in translocated regions, or that are pseudogenized are represented by colored bars on the right side of the plot.

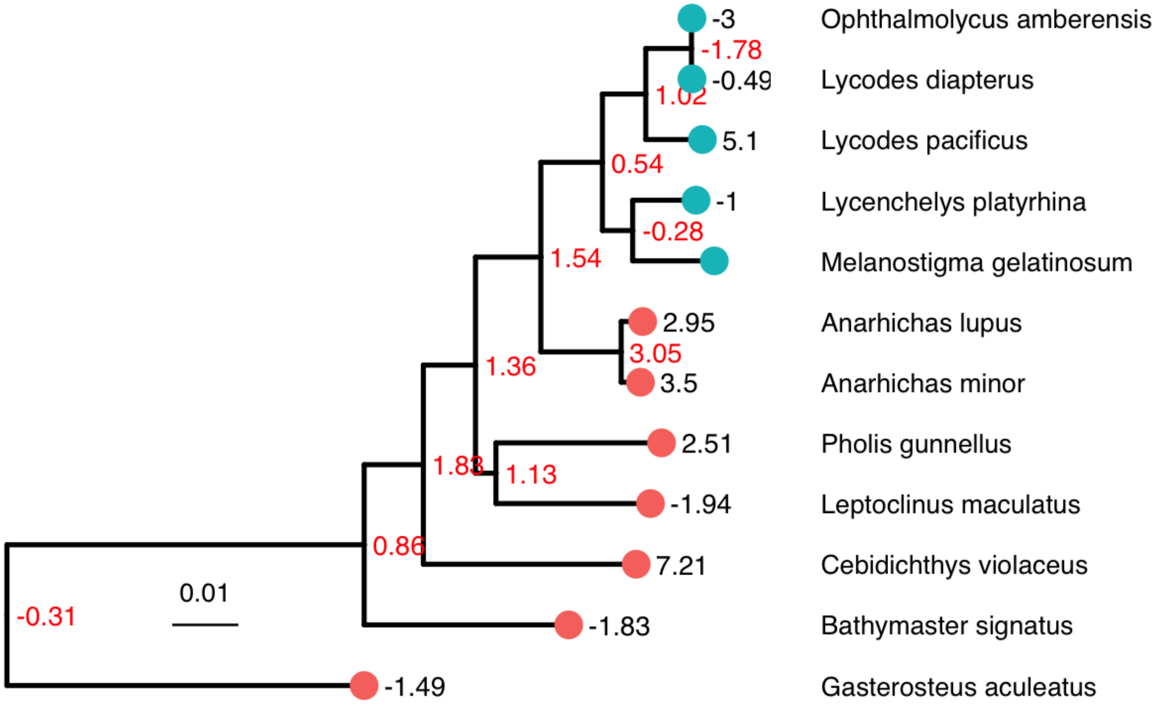


**Figure S7** | Ancestral state reconstruction of habitat thermal minima. Red values at nodes represent maximum restricted maximum likelihood predictions ancestral states in degrees Celsius. Black values represent the thermal minima of extant species at branch tips. Zoarcidae (eelpouts) branch tips are labeled with blue circles. All other zoarcoids have branch tips labeled with red circles.

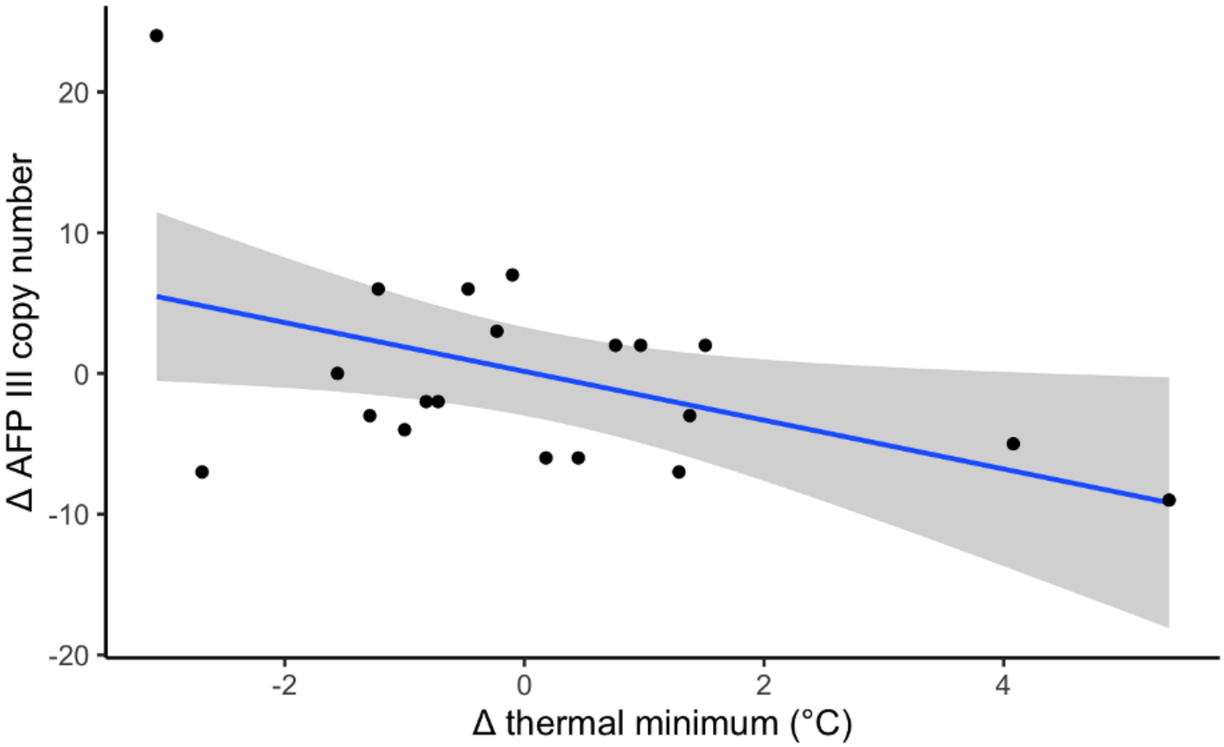


**Figure S8** | Changes in habitat thermal minima and copy number of type III antifreeze protein between nodes and descendant nodes and branches of ancestral state reconstructions. This figure serves as a visual representation of the model of convergent AFP III copy number across phylogenetically-independent variation in thermal minimum.

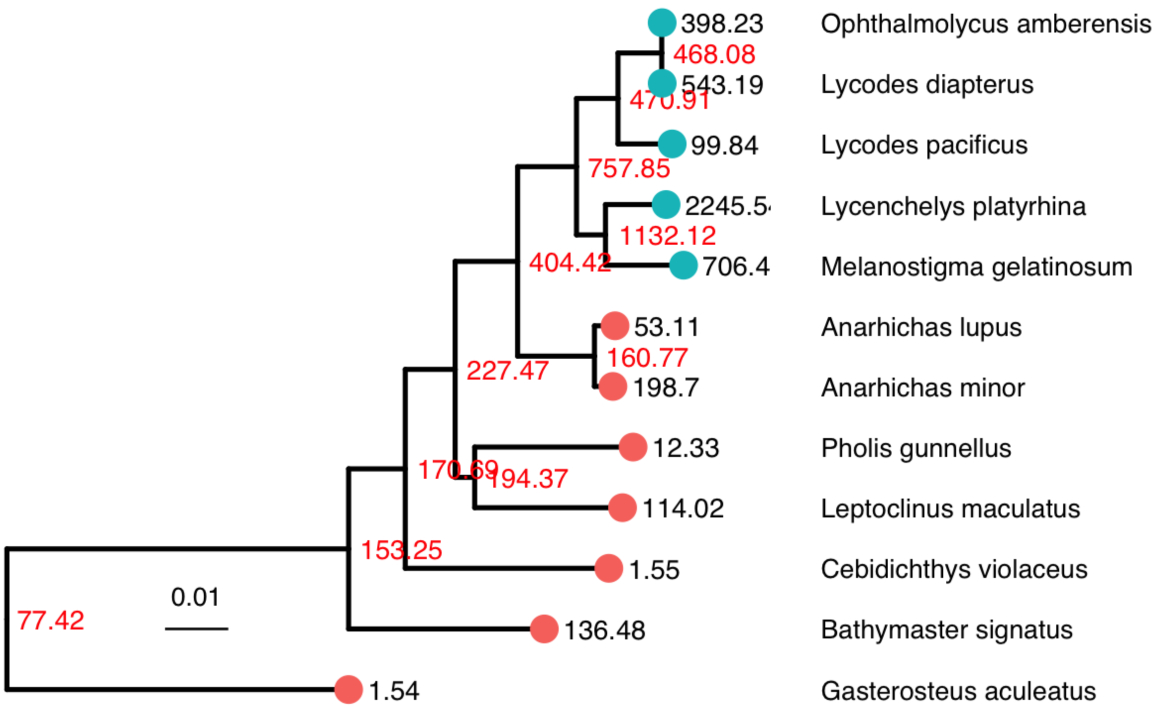


**Figure S9** | Ancestral state reconstruction of mean habitat pressure. Red values at nodes represent maximum restricted maximum likelihood predictions ancestral states in bars. Black values represent the mean habitat pressure of extant species at branch tips. Zoarcidae (eelpouts) branch tips are labeled with blue circles. All other zoarcoids have branch red circles.

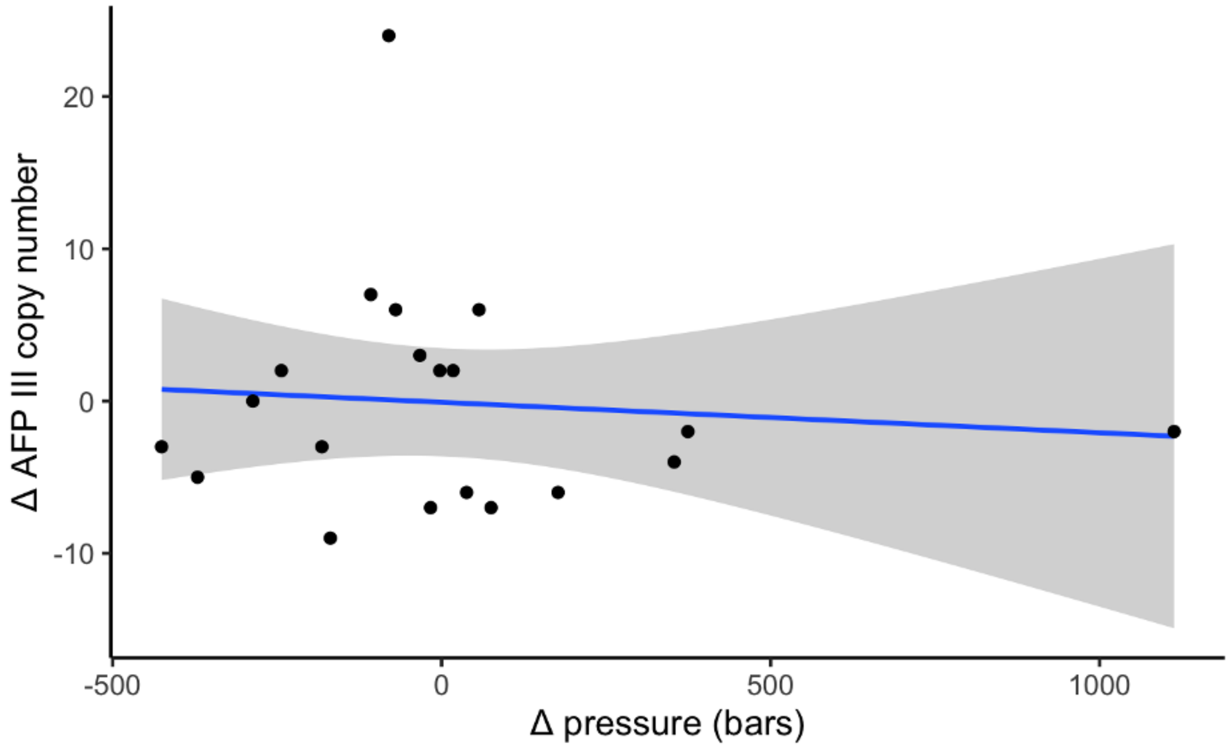


**Figure S10** | Changes in mean habitat pressure and copy number of type III antifreeze protein between nodes and descendant nodes and branches of ancestral state reconstructions. This figure serves as a visual representation of the model of convergent AFP III copy number across phylogenetically independent variation in pressure.

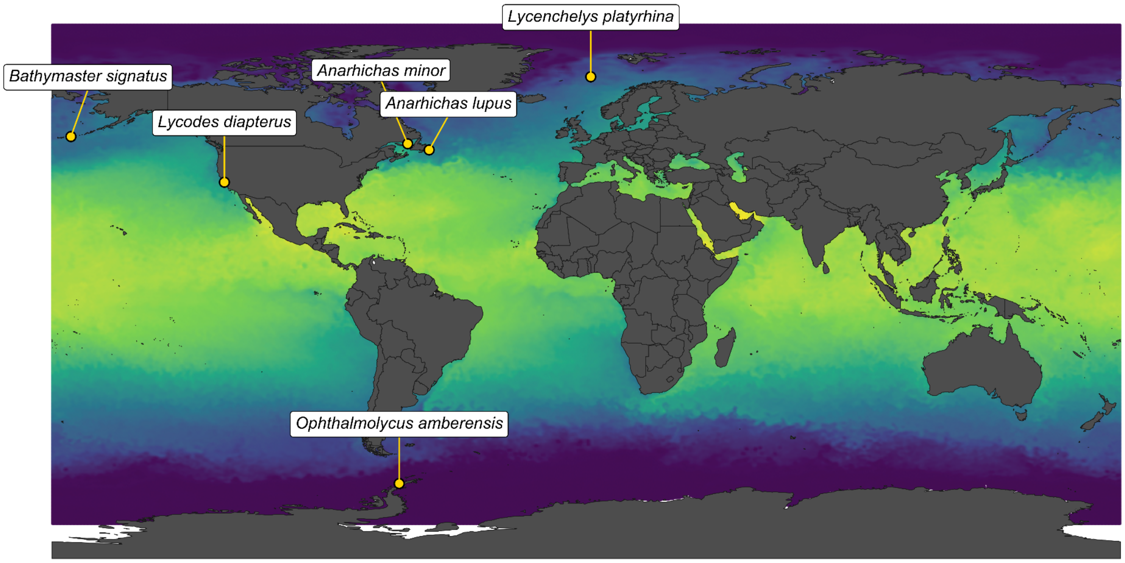


**Figure S11** | *Collection coordinates of sequenced specimens for reported assemblies and Opthalmolycus amberensis, reported by Hotaling et al. (2023).* Warm, light colors depict high sea surface temperature. Cool, dark cools depict low sea surface temperature. Mean sea surface temperature (SST, °C) recorded in August, 2020, is visualized according to color. Illustrations by SNB.

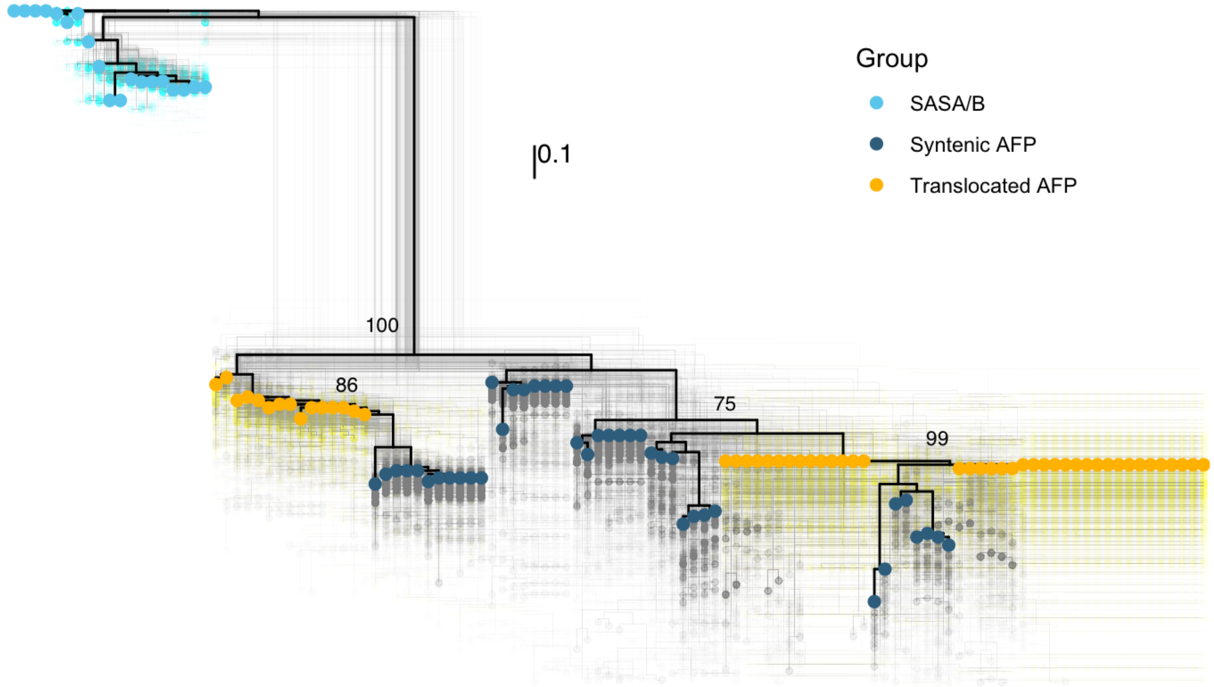


**Figure S12** | An unrooted gene tree of sasa/b and AFP III paralogs is shown. Branch tips are colored by whether they denote syntenic versus translocated AFPs or sasa/b genes. The scale bar represents nucleotide substitutions per base pair. Bootstrap support values are included to the right of all nodes that branched into clades of syntenic and translocated AFPs. The density of 1000 bootstrap trees is plotted behind the consensus gene tree.
